# Environmental co-option of chromatin potential links prenatal hypoxia to persistent cortical circuit dysfunction

**DOI:** 10.1101/2025.11.28.691133

**Authors:** Ana G. Cristancho, Donald J. Joseph, Margaret M. Cassidy, Preeti S. Chauhan, Ethan J. Gadra, Elyse C. Gadra, Donya Zarrinnegar, Bianca N. Rodriguez, Eric D. Marsh

**Affiliations:** Department of Pediatrics, Children’s Hospital of Philadelphia, Philadelphia, Pennsylvania, USA; Division of Child Neurology, Children’s Hospital of Philadelphia, Philadelphia, Pennsylvania, USA; Department of Neurology, Perelman School of Medicine at the University of Pennsylvania, Philadelphia, Pennsylvania, USA; Cell and Molecular Biology PhD Program, Perelman School of Medicine at the University of Pennsylvania, Philadelphia, Pennsylvania, USA

## Abstract

Transient prenatal hypoxia is associated with long-term neurodevelopmental impairment without overt structural brain injury, suggesting early regulatory disruptions precede detectable pathology. Whether these disruptions represent transient stress responses or redirect developmental programs remains unknown. Here, we combined single-nucleus RNA sequencing and chromatin accessibility profiling in the late-gestation mouse cortex immediately after transient hypoxia, capturing responses across more than 140,000 nuclei. Hypoxia induced a largely conserved transcriptional stress response across cell types, whereas chromatin accessibility showed more extensive, cell type–specific disruption. Developing glutamatergic neurons showed the strongest disruption of chromatin–transcription coupling, with hypoxia-responsive distal regulatory regions associated with genes governing synaptic organization and neuronal excitability. Consistent with these chromatin-associated programs, juvenile offspring displayed reduced dendritic spine density and increased intrinsic excitability in cortical neurons. These results support a model in which transient hypoxia co-opts the developing epigenome’s capacity to encode future transcriptional states before persistent circuit dysfunction emerges.

## INTRODUCTION

Transient intrauterine hypoxia is a major global contributor to long–term neurodevelopmental disability (*1, 2*). Although moderate to severe hypoxic–ischemic encephalopathy (HIE) is often accompanied by abnormalities on neonatal MRI, infants with mild HIE may have little or no MRI-detectable injury yet still develop later cognitive, behavioral, or motor impairments (*3–8*). Recent work has demonstrated that even mild neonatal hypoxia without ischemia can persistently disrupt synaptic maturation and potassium channel function in the hippocampus, yet the upstream regulatory mechanisms that establish this vulnerability at the time of injury remain unknown (*9*). This apparent “silent injury” highlights a critical gap in our understanding of how transient hypoxic exposure perturbs brain development without overt structural damage or readily detectable neuronal loss. Defining the earliest molecular consequences of hypoxia in the developing brain is therefore essential for understanding how functional deficits emerge from injuries that are not anatomically apparent at birth. Since neuronal and glial lineages undergo rapid, stage-specific differentiation during this period, identifying how hypoxia perturbs these processes at a cell type–resolved level is particularly important.

Emerging evidence indicates that the epigenome, including chromatin accessibility, histone modifications, and DNA methylation, functions as a dynamic regulator of developmental state, integrating environmental signals and shaping future cellular potential (*10–14*). During development, chromatin accessibility can precede and anticipate future transcriptional programs, including those governing synaptic gene deployment, ion channel expression, and dendritic morphogenesis (*15–20*). Joint single-cell profiling has formalized this relationship as “chromatin potential,” in which chromatin state can anticipate future transcriptional and cell-state transitions before they are evident at the RNA level (*21, 22*). In the developing brain specifically, spatial multi-omic profiling likewise demonstrates chromatin priming at neuronal and glial loci before transcriptional activation, supporting this temporal relationship during brain development (*23*). Several chromatin-modifying dioxygenases, including KDM6A, require oxygen as a cofactor, providing a direct mechanism by which transient hypoxia could perturb developmental chromatin regulation (*10*). Acute hypoxia has been shown to induce rapid and widespread chromatin remodeling, raising the possibility that even brief oxygen deprivation could disrupt the regulatory states that encode future neuronal maturation and circuit assembly before these programs are transcriptionally active (*19, 24*). While prior studies have characterized transcriptional responses to developmental brain injury, the cell type–specific epigenomic consequences of prenatal hypoxia, and whether their impact persists beyond the acute stress response, remain poorly understood (*25, 26*).

A critical unresolved question is therefore whether early changes in chromatin organization simply reflect an acute stress response or whether environmental insult can co-opt the epigenome’s normal developmental function to encode future transcriptional states and redirect that regulatory potential toward abnormal developmental outcomes. We refer to the latter process as “environmental co-option of chromatin potential.”

To address this question, we previously developed a transient prenatal hypoxia model in which pregnant mice are exposed to hypoxia late in gestation (*27*). This model produces persistent behavioral and seizure-susceptibility phenotypes despite the absence of gross neuroanatomical injury or previously detected acute cell death (*27*). Thus, it provides a model for examining early molecular responses to hypoxia while minimizing the confounding secondary cascades associated with overt tissue injury and cell loss. Given the rapid developmental transitions occurring in the fetal cortex at this time point, we reasoned that even brief hypoxia could induce lineage-specific epigenomic alterations that perturb developmental regulatory trajectories and precede persistent circuit dysfunction.

Here, we test whether transient prenatal hypoxia co-opts chromatin potential in the developing cortex by preferentially perturbing regulatory states that anticipate future transcriptional programs before their functional consequences emerge. To address this, we applied joint single-nucleus RNA and chromatin accessibility profiling (snRNA/snATAC) to the fetal cortex immediately following hypoxic exposure, enabling direct comparison of transcriptional and epigenomic responses across cell types and assessment of their regulatory coupling. By integrating these multimodal data with structural and electrophysiological analyses in juvenile offspring, we asked whether acute chromatin signatures anticipate the neuronal programs that are disrupted later in development.

## RESULTS

### Transient prenatal hypoxia largely preserves fetal cortical cell type composition

To examine how transient prenatal hypoxia perturbs molecular programs in the absence of overt injury, we profiled the fetal mouse cortex one hour after an 8–hour hypoxic exposure at embryonic day 17.5 (**Fig. 1a**). This early time point was chosen to capture the primary molecular response to hypoxia, since prior studies had suggested rapid normalization of the initial transcriptional profile (*24, 28*). This strategy allows us to isolate epigenomic and transcriptional changes that may encode risk for later dysfunction prior to those functional changes being evident.

**Fig. 1:**
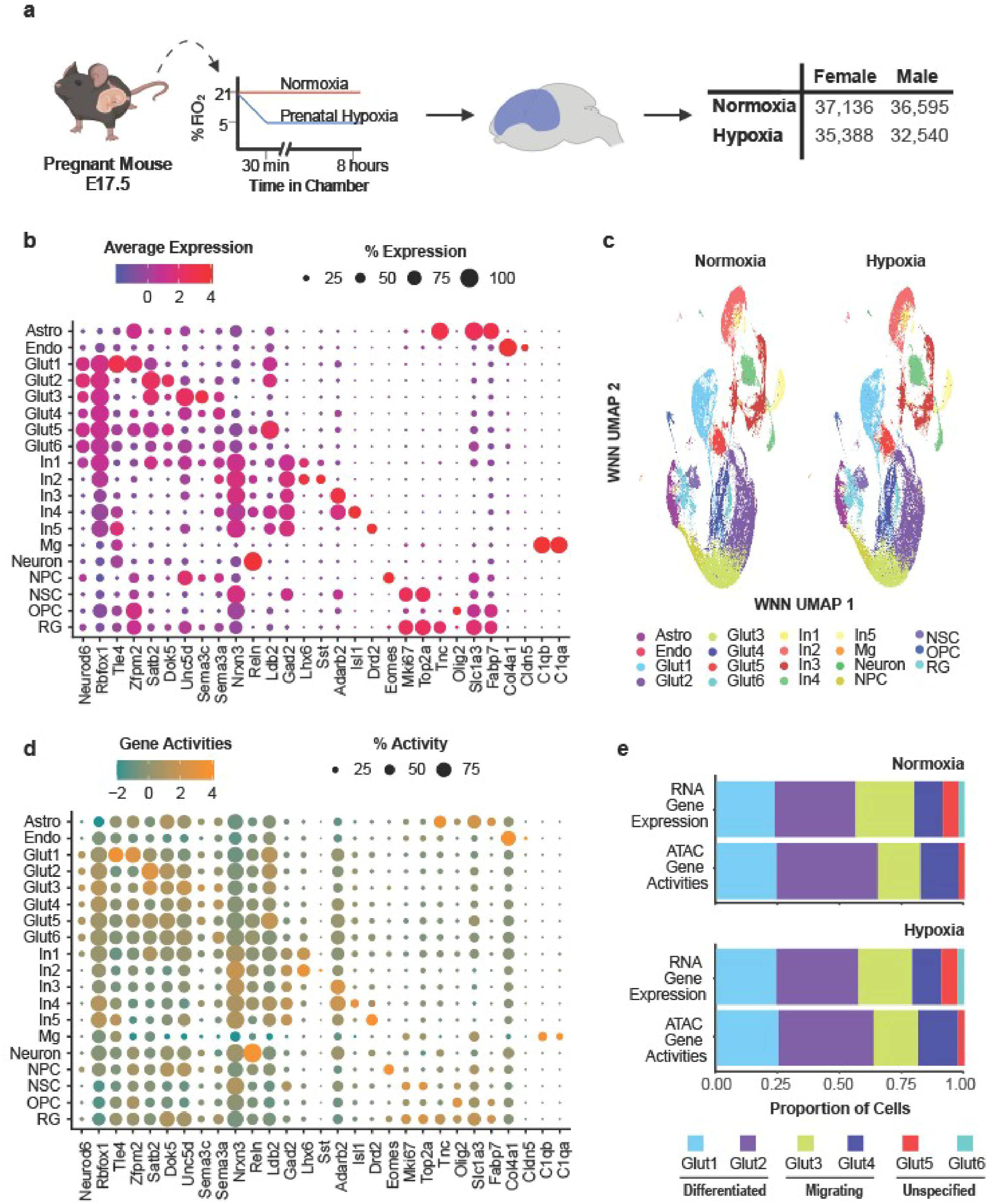
Identification and annotation of fetal cortical cell populations by joint single-nucleus multi-omics. (a) Experimental schematic of transient prenatal hypoxia at E17.5, fetal cortical collection, and joint snRNA-seq/snATAC-seq profiling; table shows nuclei retained by sex and condition. (b) Dot plot of canonical marker-gene expression across annotated cell populations; color indicates average expression and dot size indicates percentage of nuclei expressing each gene. (c) WNN UMAP shown separately for normoxia and hypoxia and colored by cell type. (d) Dot plot of chromatin-derived gene activity for the same marker genes; color indicates average gene activity and dot size indicates percentage of nuclei with detectable activity. (e) Relative representation of glutamatergic subclasses defined by RNA expression or ATAC gene activity, shown separately for normoxia and hypoxia.

We sequenced 163,240 nuclei from 16 biological samples evenly distributed by sex and condition using the 10× Genomics Joint Multi-ome platform. After stringent quality–control filtering (**Table S1**), 141,659 nuclei were retained, with a median of 2,202 genes detected per nucleus and 8,752 high–quality ATAC fragments per nucleus, providing a substantial cohort to detect subtle regulatory changes independent of cell number or developmental state. Before integration, RNA-derived representations showed little segregation by experimental condition, whereas ATAC-derived representations showed pronounced separation between normoxia and hypoxia (**Fig. S1 & S2**). Comparable nucleosomal distribution and transcription start site (TSS) enrichment between conditions argued against global differences in sequencing quality as the sole explanation for the condition-dependent ATAC signal (**Fig. S3**). To facilitate joint integration and consistent cell-type annotation across conditions, we applied Harmony to the ATAC LSI representation before weighted nearest neighbor integration (**Fig. S4**) (*29*). Harmony-adjusted embeddings were used for integration and visualization only and were not used for differential accessibility analyses.

An iterative approach based on RNA signatures defining cell classes (glutamatergic neurons, interneurons, proliferating cells, and glia) and then subclasses, identified expected cortical populations, including glutamatergic neurons, interneurons, neural progenitors, proliferative cells, astrocytes, oligodendrocyte progenitor cells, microglia, and endothelial cells (**Fig. 1b–c; Fig. S5a**) (*30–35*). At this developmental stage, neuronal subclasses represent transitional states rather than terminal identities; therefore, subclass nomenclature reflects transcriptional similarity to mature populations rather than definitive fate commitment (**Table S2**) (*33, 36*). Cell type proportions were largely preserved after hypoxia (**Fig. S5b–c**), indicating that the acute molecular response occurred without major reorganization of cortical cellular composition. The exception was an increase in endothelial cells, consistent with known hypoxia–driven angiogenic signaling(*37*). Progenitor populations exhibited expected proliferative signatures, but no differences in cell–cycle score distributions were detected between normoxia and hypoxia (**Fig. S5d**).

Chromatin–derived gene activity scores, defined by quantification of open chromatin within the gene body as a proxy for gene expression, did not fully resolve finer neuronal subtypes, particularly among migratory glutamatergic neurons (**Fig. 1d; Table S3**) (*31, 38*). This limited resolution likely reflects both the inherent properties of gene activity scores and developmental asynchrony between chromatin accessibility and transcriptional identity in maturing excitatory neurons. Importantly, this pattern was observed in both normoxic and hypoxic samples (**Fig. 1e**), indicating that it reflects developmental chromatin dynamics rather than injury per se. Critically, this baseline dissociation, in which chromatin state encodes future transcriptional potential not yet reflected in current gene expression, is precisely the developmental mechanism that hypoxia appears to co-opt, making its characterization under normoxic conditions essential context for interpreting later hypoxia-induced discordance.

Together, these data show that fetal cortical cell type composition is largely preserved immediately after transient prenatal hypoxia, despite condition-dependent differences in the ATAC-derived representation. This prompted us to examine transcriptional and chromatin responses within individual cortical populations.

### A conserved transcriptional response to hypoxia spans fetal cortical cell types

We next examined hypoxia-induced transcriptional changes across cell types. Principal component analysis of differential expression fold changes revealed strong alignment of response vectors across neuronal, glial, and progenitor populations (**Fig. 2a**), indicating that when genes respond to hypoxia, they tend to do so in a shared direction regardless of lineage.

**Fig. 2:**
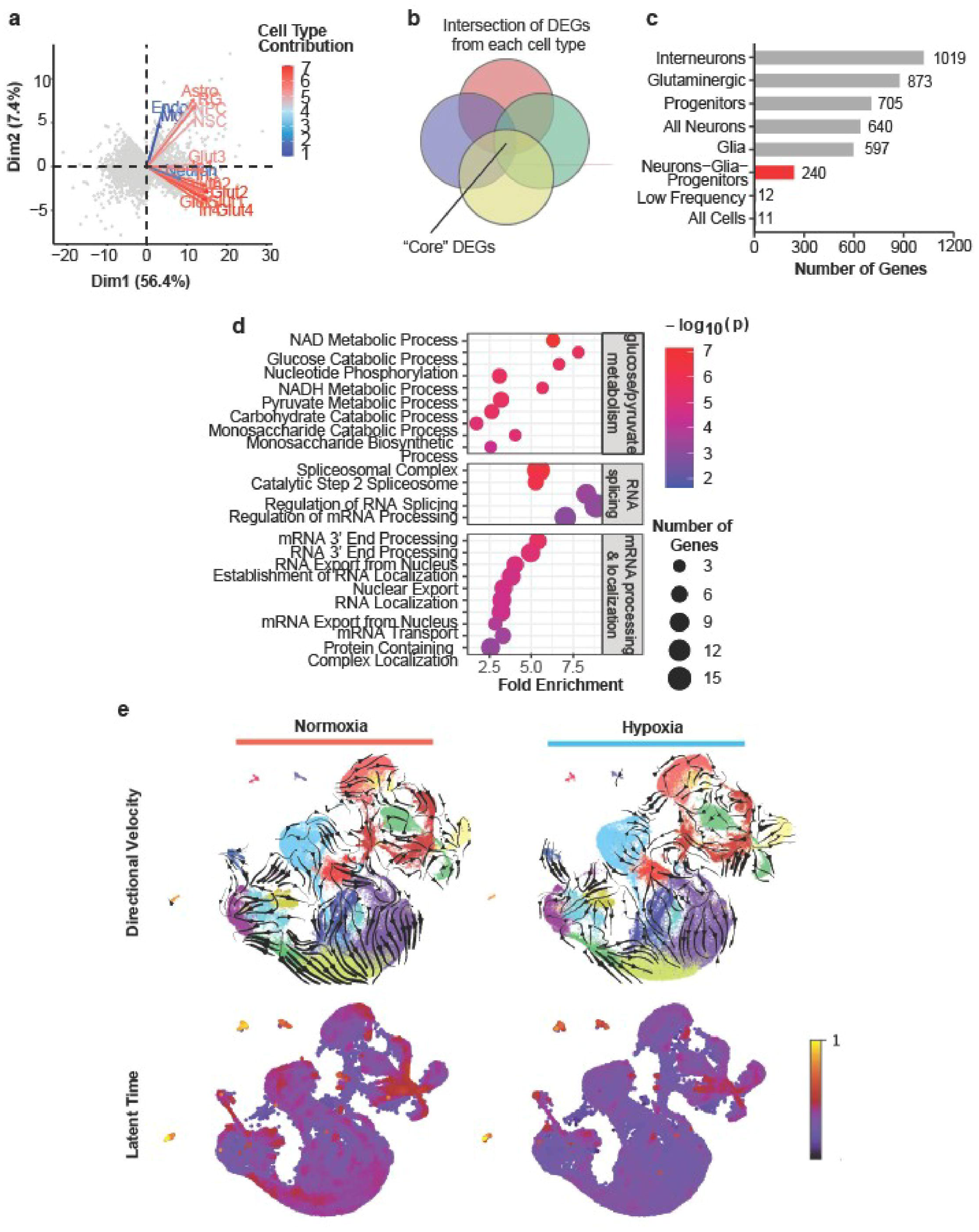
A conserved transcriptional response to prenatal hypoxia spans fetal cortical cell types. (a) Principal component analysis of hypoxia-associated differential-expression fold changes across cortical cell populations; vectors indicate cell type contributions to the principal-component space. (b) Schematic defining “core” hypoxia-responsive genes as the intersection of differentially expressed genes across cell types. (c) Number of overlapping hypoxia-responsive genes across increasingly broad cell-type groupings; the 240 genes shared across abundantly represented neuronal, glial, and progenitor populations are highlighted in red. (d) Gene ontology enrichment of core hypoxia-responsive genes. Dot position indicates fold enrichment, color indicates adjusted −log10(p), and size indicates number of genes. (e) RNA velocity vector fields and latent-time projections for normoxia and hypoxia.

Intersecting differentially expressed genes (adjusted *P* < 0.05, |log₂ fold change| > 0.25) from abundantly represented populations (>1,000 cells per type) identified 240 core hypoxia-responsive genes shared across neurons, glia, and progenitors (**Fig. 2b–c**). The inclusion of low-frequency populations markedly reduced overlap, likely due to limited statistical power, and these populations were excluded from shared signature analyses.

Gene ontology (GO) enrichment analysis revealed that core genes were enriched for pathways related to energy metabolism and RNA binding and processing (**Fig. 2d**), consistent with a conserved response to energetic stress(*39*). Given the prominence of RNA regulatory terms, we hypothesized that hypoxia altered RNA kinetic dynamics. RNA velocity analysis using a dynamical model showed disorganized velocity fields and latent time distributions under hypoxia (**Fig. 2e**). Kinetic parameter estimation indicated globally reduced transcription and degradation rates without changes in splicing rates (**Fig. S6a–c**) (*40*).

These findings indicate that hypoxia induces a broadly conserved transcriptional response across cortical cell types, accompanied by altered RNA kinetic dynamics rather than a coherent acceleration or arrest of differentiation.

### Cell type-selective transcriptional responses converge on shared regulatory processes

To identify lineage-specific transcriptional responses beyond the shared hypoxia program, we analyzed cell type–selective differentially expressed genes (DEGs) after excluding the 240 core hypoxia-responsive genes shared by neurons, glia, and progenitors. This approach allowed us to separate transcriptional changes more unique to each population from the global stress responses driven by energetic deprivation. Because individual genes could appear in multiple cell type–selective sets, these signatures reflect preferential rather than exclusive association with a given lineage.

To assess potential clinical relevance, mouse DEGs were mapped to human orthologs and analyzed using the DisGeNET database (*41*). Both core and cell type–selective gene sets showed significant enrichment for terms associated with neurodevelopmental and neurological disorders, including intellectual disability, epilepsy, motor dysfunction, and behavioral phenotypes, across neuronal and non-neuronal populations (**Fig. 3a**). Importantly, size-matched random gene lists did not show comparable enrichment for neurodevelopmental phenotypes, supporting the specificity of these associations. Notably, enrichment for neurodevelopmental and neurological phenotypes recurred across cell types despite substantial differences in the underlying hypoxia-responsive genes, suggesting that distinct lineage-specific transcriptional responses may converge on biological processes relevant to shared neurodevelopmental outcomes.

**Fig. 3:**
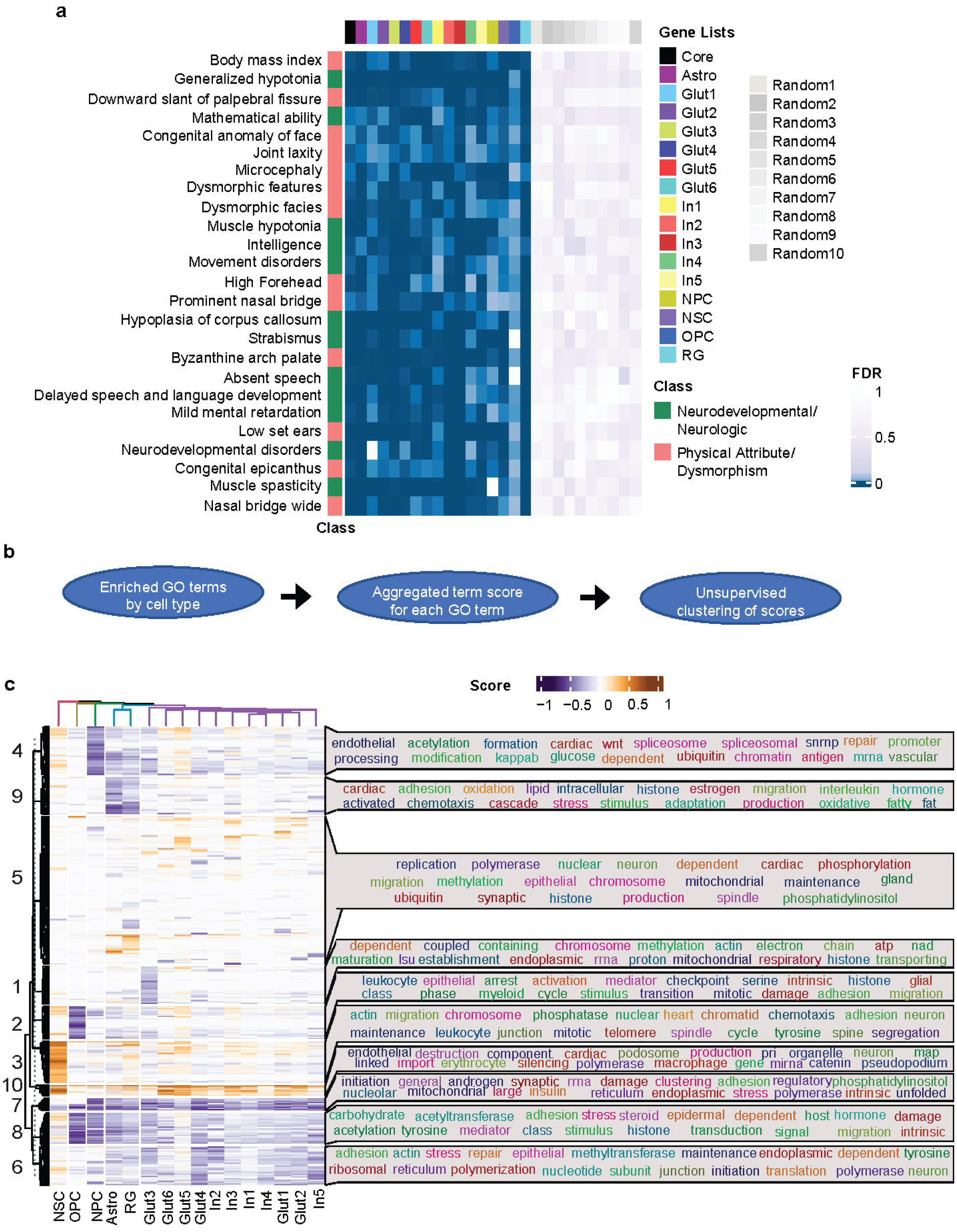
Cell type-selective transcriptional responses converge on shared regulatory and developmental processes. (a) DisGeNET phenotype enrichment among core and cell type-selective hypoxia-responsive gene sets. Heatmap color represents FDR; phenotype classes are indicated at left. Size-matched random gene sets are shown at right as controls. (b) Schematic of the workflow used to aggregate and cluster GO-term enrichment across cell types. (c) Heatmap of aggregated GO-term enrichment scores across cell types, with unsupervised clustering into 10 pathway clusters; word clouds summarize recurrent biological terms within each cluster.

Recognizing the potential for bias and overinterpretation in disease-association analyses of large gene sets, we next examined functional convergence independently of disease annotation. GO enrichment was performed for each cell type-selective DEG set, and enrichment scores were aggregated and subjected to unsupervised clustering (**Fig. 3b–c**). This approach assessed whether distinct cell type-selective gene programs converged on shared biological processes.

Clustering revealed a clear segregation between neuronal populations and glial or progenitor populations, consistent with transcriptional PCA results (**Fig. 2a**). Neural progenitors were preferentially enriched for pathways related to cell division, DNA replication, and mitotic regulation, suggesting hypoxia-induced perturbation of proliferative programs (*cluster 3*). Oligodendrocyte progenitor cells showed enrichment for pathways associated with cellular stress, damage responses, and growth arrest, consistent with the known sensitivity of these cells to hypoxic injury (*cluster 2*) (*42, 43*). Despite the heterogeneity of differentially expressed genes across populations, a striking convergence emerged at the pathway level. Across lineages, hypoxia-responsive gene sets repeatedly implicated transcriptional regulation, chromatin organization, and epigenetic control, indicating that alterations in gene regulatory machinery represent a unifying consequence of prenatal hypoxia.

Together, these findings indicate that prenatal hypoxia elicits distinct cell type-selective transcriptional responses that nevertheless converge on shared regulatory and developmental processes. The repeated implication of chromatin organization and epigenetic regulation across lineages suggested that disruption of developmental regulatory architecture may represent a common feature of the hypoxic response, motivating direct examination of chromatin accessibility.

### Hypoxia preferentially disrupts chromatin organization in developing glutamatergic neurons

Given the relatively conserved transcriptional response to hypoxia across cortical cell types, we next asked whether prenatal hypoxia produced a more differentiated, lineage-specific response at the level of chromatin accessibility. Because chromatin state can precede measurable transcriptional output during development, accessibility profiling provides a means to identify regulatory perturbations that may not yet be evident at the RNA level (*21, 44*).

To assess global effects on developmental progression, we first performed pseudotime ordering of transcriptomic trajectories across major cortical lineages relative to the progenitor NSC population. Consistent with the modest transcriptional differences observed at this early time point, transcriptomic pseudotime analysis revealed only subtle shifts between normoxic and hypoxic samples across neural progenitors, neuronal populations, and glial lineages (**Fig. S7**). These findings indicate that transient prenatal hypoxia does not grossly alter RNA-defined developmental trajectories immediately following exposure.

Whereas transcriptomic trajectories remained largely overlapping between conditions, accessibility-based trajectories showed widespread displacement after hypoxia(**Fig. 4a**). Together with the pronounced condition-dependent separation of ATAC-derived representations, these findings reveal a substantially greater reorganization of chromatin-defined state than of transcriptionally defined identity at this acute time point.

**Fig. 4:**
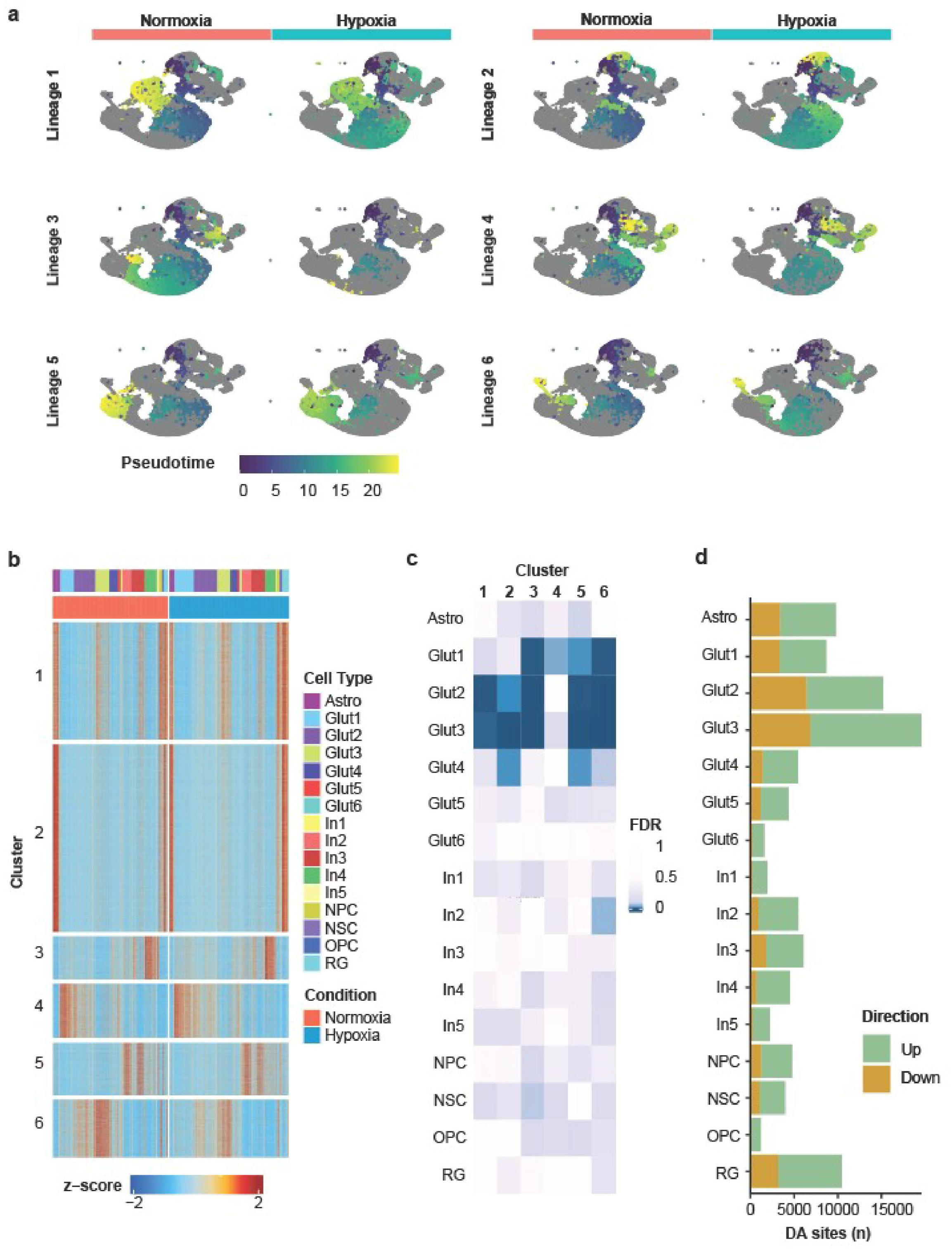
Prenatal hypoxia disproportionately disrupts chromatin regulatory architecture in developing glutamatergic neurons. (a) Chromatin accessibility-based pseudotime projected onto WNN UMAP embeddings for six cortical lineages, shown separately for normoxia and hypoxia. Cells within each lineage are colored by pseudotime; cells outside the indicated lineage are shown in gray. Pseudotime distributions differed between normoxia and hypoxia across all six lineages (two-sample Kolmogorov–Smirnov test, p < 2.2 × 10⁻¹⁶). (b) Peak-to-gene linkage heatmap across fetal cortical cell types and conditions, partitioned into six linkage clusters. Heatmap color represents z-scored peak-to-gene linkage signal; annotations indicate cell type and condition. (c) FDR of permutation-based comparisons of peak-to-gene linkage density between normoxia and hypoxia for each cell type and linkage cluster. (d) Number of hypoxia-associated differentially accessible sites by cell type, separated into regions with increased or decreased accessibility following hypoxia.

To interrogate these epigenomic changes at higher resolution, we recalculated chromatin accessibility peaks independently within each cell type using ArchR’s fixed-width iterative overlap merging approach(*45*), identifying a total of 604,493 accessible regions across all populations (**Fig. S8**). Under baseline conditions, peaks-to-gene linkage analysis revealed strong and expected cell type specificity, with regulatory interactions aligning closely with transcriptionally defined identities (**Fig. 4b**), consistent with prior reports in the developing and adult cortex(*35, 46*).

Permutation-based comparisons of peak-to-gene linkage density between conditions revealed marked cell type–specific differences. Among all analyzed populations, glutamatergic neurons exhibited the greatest hypoxia-associated disruption of peaks-to-gene linkage, substantially exceeding that observed in interneurons, glial populations, or progenitor cells (**Fig. 4c**; **Fig. S9**). Thus, the regulatory coupling between chromatin accessibility and transcription is disproportionately disrupted in developing excitatory neurons following hypoxia.

Consistent with this observation, differential accessibility analysis revealed that the glutamatergic neuron subtypes Glut2 and Glut3 harbored the highest number of hypoxia-responsive chromatin sites (**Fig. 4d**). Hypoxia-responsive regions showed limited overlap across cell types (**Fig. S10**). Hypoxia-downregulated sites were preferentially located in distal enhancer regions, whereas upregulated sites were more commonly found near promoter-proximal regions (**Fig. S11)**. Because regulatory elements play a central role in establishing lineage-specific transcriptional programs during cortical development, their preferential disruption in glutamatergic neurons suggests that hypoxia targets regulatory architecture involved in subsequent neuronal maturation (*15, 47*).

Importantly, although interneurons and glial populations also exhibited hypoxia-associated accessibility changes, these changes were fewer in number and produced significantly less disruption of regulatory architecture compared with glutamatergic neurons (**Fig. 4d**). This pattern parallels developmental studies showing that excitatory projection neurons undergo extensive chromatin reconfiguration during late gestation as they transition from migratory to early post-mitotic states, potentially rendering them uniquely sensitive to perturbations in oxygen-dependent epigenetic processes (*18, 19*).

These data demonstrate that prenatal hypoxia disproportionately destabilizes the chromatin regulatory architecture of developing glutamatergic neurons, producing greater disruption of chromatin–transcription coupling than observed in other cortical lineages. This vulnerability emerges despite largely preserved cellular composition and modest changes in RNA-defined developmental trajectories, identifying chromatin dysregulation as the dominant lineage-specific molecular signature of the acute hypoxic response.

### Early chromatin accessibility signatures anticipate persistent structural and functional dysfunction in glutamatergic neurons

We next asked whether the acute epigenomic changes observed after prenatal hypoxia were informative of long-term neuronal outcomes. We focused on Glut1 (corticothalamic) and Glut2 (callosal projection) neurons, as these subtypes represent early maturing excitatory populations that are actively undergoing circuit integration at this developmental stage.

Gene ontology analysis of differentially expressed genes in Glut1 and Glut2 neurons revealed enrichment primarily for pathways related to translation and oxygen sensing (**Fig. 5a**), consistent with an acute metabolic response to hypoxic stress. In contrast, gene ontology analysis of genes neighboring differentially accessible chromatin regions, identified independently of transcriptional changes, uncovered a striking enrichment for pathways involved in synaptic organization, synaptic plasticity, and long-term potentiation (**Fig. 5b**). These functional categories are directly relevant to later neuronal connectivity and circuit function yet were not apparent from transcriptional analysis at the fetal time point. Thus, acute transcription predominantly captured the immediate response to hypoxia, whereas chromatin accessibility implicated neuronal programs with functions that emerge later in circuit maturation. These findings are consistent with the concept of chromatin potential, in which accessibility states can reveal prospective developmental programs before they are evident in contemporaneous transcription (*21, 22*).

**Fig. 5:**
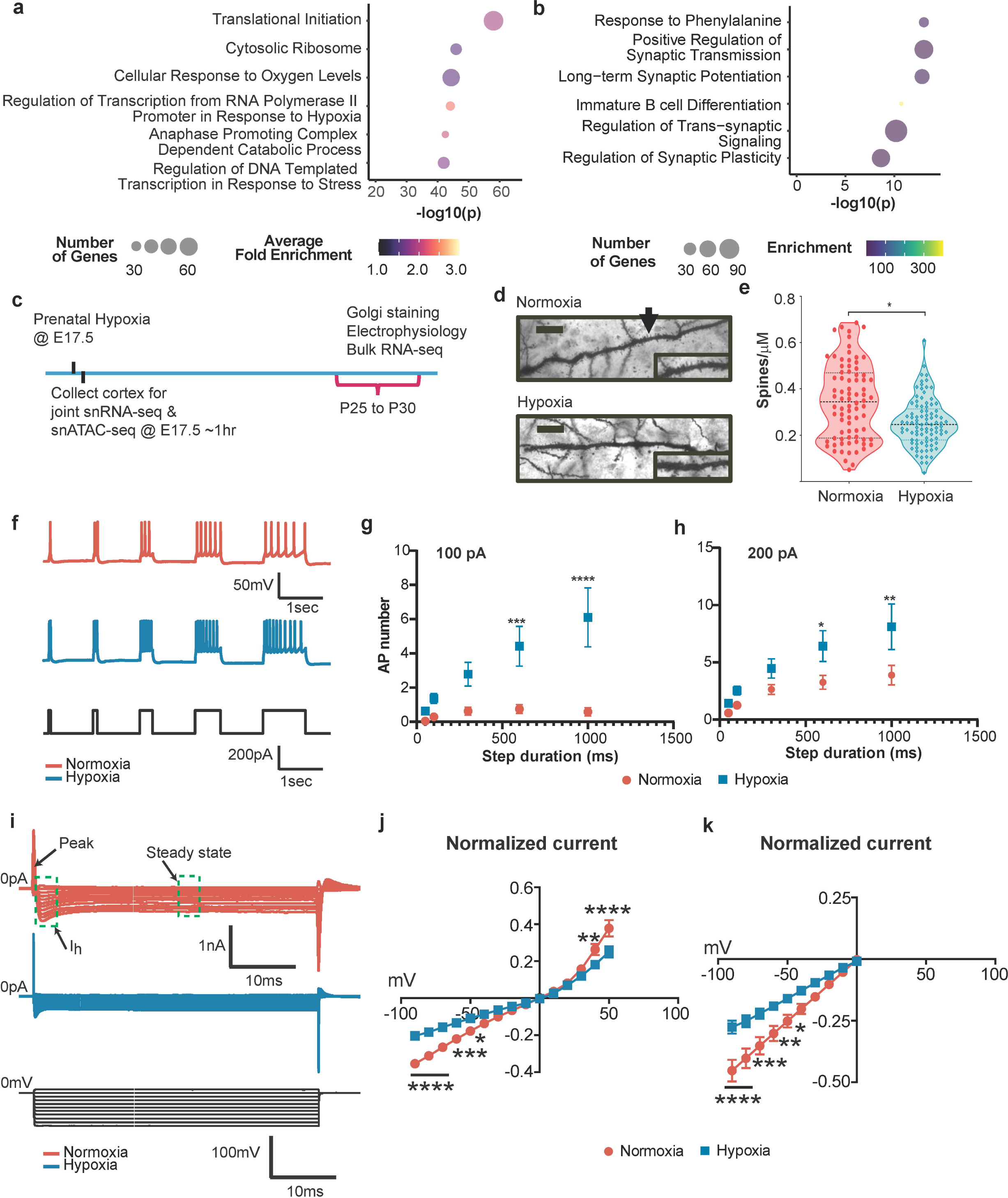
**Fetal chromatin accessibility signatures anticipate persistent structural and electrophysiological abnormalities in glutamatergic neurons.**(a) Gene ontology enrichment of differentially expressed genes in Glut1 and Glut2 neurons immediately following prenatal hypoxia. Dot position indicates calculated combined −log10(p), size indicates number of genes, and color indicates average fold enrichment. (b) Gene ontology enrichment of genes neighboring hypoxia-associated differentially accessible regions in Glut1 and Glut2 neurons. Dot position indicates adjusted −log10(p), size indicates number of genes, and color indicates enrichment. (c) Experimental timeline for fetal multi-omic profiling and juvenile Golgi staining, electrophysiology, and bulk RNA-seq. (d) Representative images of golgi staining quantified in (e); nested two tailed t-test, mean µm ± SD & cells/animals; normoxia: 0.34 ± 0.16 & 80/3, hypoxia 0.25 ± 0.10 & 93/3, *p =* 0.0219). Scale bar = 20 µm) Increased membrane excitability of Pyramidal cells by prenatal hypoxia. (f) Representative traces of pyramidal cell responses to 200pA step current injections in normoxia (coral) and hypoxia (turquoise) given at indicated step durations; pulses as stimulus waveform (Bottom). (g, h) Summary of data for action potential number elicited at different step durations for 100pA (g) and 200pA (h) injected current steps. Normoxia, Hypoxia: *n=*24/8, 19/6, slices/mice; F(1, 205) = 40.9 for 100pA plot, F (1, 205) = 18.4 for 200pA plot; 100pA/600ms normoxia=0.75±0.26 vs. hypoxia= 4.42±1.16; 200pA/600ms normoxia =3.25±0.61 vs. hypoxia = 6.42±1.35 and 100pA/1000ms normoxia=0.58±0.25 vs. hypoxia= 6.10±1.72 and 200pA/1000ms normoxia=3.87±0.85 vs. hypoxia = 8.10±1.99. (i) Representative traces of the current-voltage (I-V) relationship recorded from PCs in normoxia (coral) and hypoxia (turquoise) in the voltage clamp mode. (j, k) Plots of the I-V relationship for the steady state or potassium current (j; normoxia, hypoxia: *n=* 25/24, 25/19, slices/mice; F(1, 710) = 21.3) and Ih current (k; normoxia, hypoxia: *n=* 25/24, 25/19, slices/mice; F(1, 520) = 48.8, *p* < 0.0001) components. Electrophysiology data are given as mean±SEM and analyzed by Two-way ANOVA followed by Šídák’s multiple comparisons test: \**p*<0.05, \*\**p*<0.01, \*\*\**p*<0.001, \*\*\*\**p*<0.0001.

We therefore asked whether the neuronal programs implicated by fetal chromatin accessibility were reflected in later cellular phenotypes (**Fig. 5c**). In prenatal exposed juvenile animals, Golgi staining revealed a significant reduction in dendritic spine density in layer 5/6 pyramidal neurons of the anterior cingulate cortex (**Fig. 5d–e**), a region implicated in behavioral phenotypes previously observed in this model (*27, 48*).

Reduced spine density is a well-established correlate of impaired synaptic connectivity and plasticity in neurodevelopmental disorders, so we next assessed whether these structural abnormalities were accompanied by functional deficits (*49–51*). Whole-cell patch-clamp recordings from pyramidal neurons revealed increased action potential firing, reduced sustained potassium currents, diminished I_h_ currents, and altered passive membrane properties in animals exposed to prenatal hypoxia (**Fig. 5f–k**; **Fig. S12**). These changes indicate persistent neuronal hyperexcitability and impaired intrinsic membrane regulation, consistent with disruption of synaptic and ion channel–associated pathways identified through chromatin accessibility analysis. Notably, the later structural and electrophysiological phenotypes aligned with the synaptic and excitability programs identified by fetal chromatin accessibility but not with the dominant pathways identified by acute fetal transcription.

Together, these findings show that acute chromatin accessibility captures developmental programs that precede and anticipate persistent excitatory-neuron dysfunction more clearly than the contemporaneous transcriptional response.

### Early epigenomic dysregulation at potassium channel loci links fetal hypoxia to delayed excitability phenotypes

The co-option framework predicts that hypoxia should alter chromatin accessibility at loci that encode future functional programs before those genes become transcriptionally active. To test this prediction directly, we examined voltage-gated potassium channel genes, whose reduced currents were a prominent electrophysiological phenotype in juvenile animals (**Fig. 5; Fig. S12**). Analysis of snRNA–seq data revealed that most K⁺ channel genes were minimally expressed in the fetal cortex at embryonic day 17.5, with no significant differences between normoxic and hypoxic conditions (**Fig. 6a**). To place this observation in a developmental context, we examined single–cell gene expression data from the adult cortex using publicly available datasets, which demonstrated robust expression of multiple voltage–gated potassium channel genes later in life (**Fig. 6b**) (*35*). These findings indicate that a lack of fetal transcription does not preclude later functional relevance and suggest that regulatory mechanisms acting prior to gene expression may shape subsequent ion–channel deployment.

**Fig. 6:**
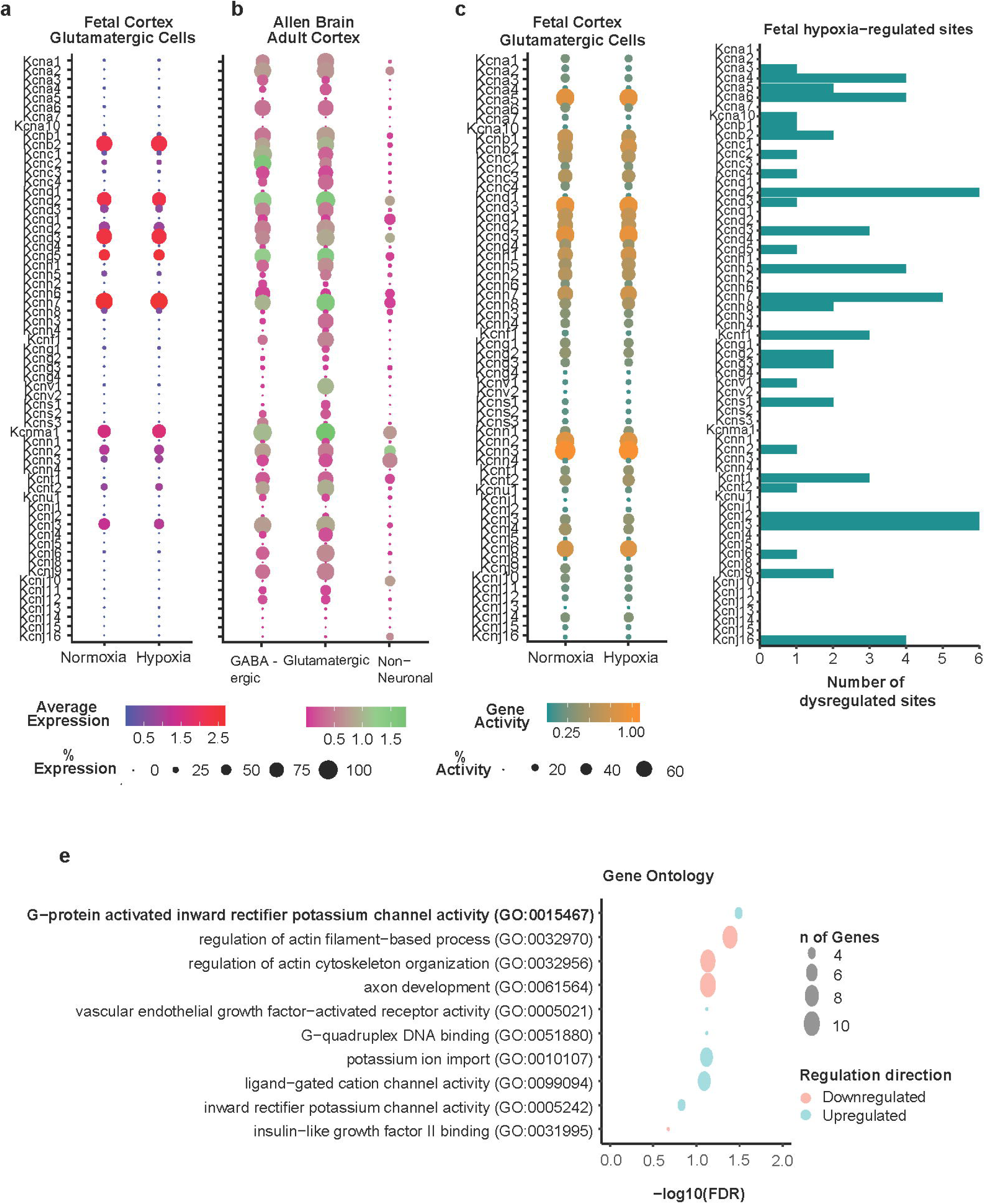
Early chromatin dysregulation at potassium channel loci precedes later transcriptional and functional abnormalities. (a) Expression of voltage-gated potassium channel genes in fetal glutamatergic neurons following normoxia or prenatal hypoxia. Color indicates average expression and dot size indicates percentage of nuclei expressing each gene. (b) Expression of potassium channel genes in GABAergic, glutamatergic, and non-neuronal populations of the adult mouse cortex using Allen Brain Atlas data. (c) Chromatin-derived gene activity at potassium channel loci in fetal glutamatergic neurons following normoxia or hypoxia. Color indicates gene activity and dot size indicates percentage of nuclei with detectable activity. (d) Number of hypoxia-associated differentially accessible sites neighboring each potassium channel gene. Enrichment relative to randomly selected genes is shown in Supplementary Fig. 13. (e) Gene ontology enrichment of differentially expressed genes from juvenile anterior cingulate cortex bulk RNA-seq. Dot position indicates −log10(FDR), size indicates number of genes, and color indicates direction of differential expression.

We next assessed chromatin accessibility at potassium channel loci in the fetal cortex. Despite minimal fetal transcription, many voltage-gated potassium channel loci were already accessible in fetal glutamatergic neurons (**Fig. 6c**), consistent with a permissive or primed regulatory state. Strikingly, approximately 50% of these loci were associated with neighboring hypoxia-responsive differentially accessible regions, significantly more than expected for randomly selected genes (**Fig. 6d; Fig. S13;** Chi-square \*\*\*\**p*<0.0001). These differentially accessible regions overlapped with previously characterized enhancer elements associated with neuronal subtype specificity, supporting their potential regulatory relevance(*46*).

This pattern suggests that prenatal hypoxia does not immediately alter potassium channel transcription, but instead reshapes the regulatory landscape surrounding these genes, before their robust developmental expression. Such epigenomic priming has been shown in developmental systems to precede transcriptional activation and to encode future gene expression programs, particularly for genes involved in neuronal excitability and synaptic function (*21, 22, 52*). To determine whether these early chromatin alterations were associated with delayed transcriptional outcomes, we performed bulk RNA–sequencing of the anterior cingulate cortex from juvenile mice exposed to prenatal hypoxia. Pathway analysis revealed that potassium channel–related gene sets were among the most dysregulated pathways at this later time point (**Fig. 6e**). These transcriptional changes align closely with electrophysiological findings of reduced sustained K⁺ currents and altered intrinsic membrane properties in the same region, providing a developmental link between early regulatory disruption and later molecular and functional abnormalities.

Together, these data define a coherent developmental sequence from acute regulatory disruption to delayed molecular and physiological dysfunction: prenatal hypoxia alters chromatin accessibility at ion channel loci before their robust fetal expression, followed later by dysregulation of potassium channel pathways and impaired neuronal excitability. This temporal sequence provides a concrete example of environmental co-option of chromatin potential, in which disruption of a prospective developmental regulatory program precedes the phenotype associated with that program.

## DISCUSSION

Transient prenatal hypoxia is strongly associated with later neurodevelopmental disability, even in the absence of detectable structural injury on neonatal imaging (*3–8*). Our findings provide a regulatory framework for this ’silent’ injury: brief late-gestation hypoxia disproportionately perturbs chromatin states that normally anticipate future transcriptional programs, despite largely preserved cellular composition and only modest acute changes in RNA-defined developmental trajectories. By integrating single-nucleus transcriptomic and epigenomic profiling with structural and electrophysiological analyses across development, we show that these early regulatory signatures are associated with later alterations in neuronal connectivity and intrinsic excitability.

Three principal findings support this conclusion. First, prenatal hypoxia elicits a largely conserved transcriptional response across neurons, glia, and progenitor populations, dominated by pathways related to energy metabolism and RNA regulation, consistent with a broad cellular response to energetic stress (**Fig. 2**). Cell type-selective transcriptional responses nevertheless converge on shared regulatory and developmental processes, including chromatin organization and epigenetic control (**Fig. 3**). Second, chromatin accessibility shows a more differentiated response across cell types, with pronounced lineage-specific effects concentrated in developing glutamatergic neurons (**Fig. 4**). Third, the biological programs implicated by fetal chromatin accessibility align closely with later structural, transcriptional, and electrophysiological abnormalities that were not prominent in the contemporaneous fetal transcriptional response (**Figs. 5 & 6**). Together, these findings suggest that chromatin accessibility captures prospective developmental information about the consequences of prenatal injury that is not evident from acute gene expression alone.

At the transcriptional level, hypoxia induced a conserved response dominated by energy metabolism and RNA processing pathways, with disorganized RNA velocity dynamics and altered kinetic parameters rather than a coherent directional shift in differentiation (**Fig. 2**). These transcriptional changes occurred in the context of preserved cell type proportions and cell-cycle distributions, indicating that the acute response was not accompanied by major reorganization of cortical cellular composition. In marked contrast, chromatin accessibility trajectories showed pronounced condition-dependent differences across cortical lineages (**Fig. 4**), with the most pronounced effects in glutamatergic neurons, including substantial disruption of peak-to-gene linkage and preferential alteration of distal regulatory elements. Because chromatin accessibility can precede transcriptional output during development, this divergence is particularly relevant in the fetal cortex, where regulatory programs required for later circuit maturation may already be established before they are fully transcriptionally deployed.

The dissociation between transcriptional readouts and chromatin-based anticipation of neuronal function (**Figs. 5 and 6**) provides the most direct evidence for environmental co-option in this study. Fetal transcription predominantly reflects the immediate cellular response to hypoxia, metabolic adaptation, and oxygen sensing, while chromatin accessibility at the same time point implicates synaptic plasticity and excitability programs whose functions become increasingly important during subsequent neuronal maturation. These pathways were not prominent in fetal transcription, but corresponding structural and physiological abnormalities emerged later, including reduced dendritic spine density, increased intrinsic excitability, and altered membrane currents. Hypoxia may therefore co-opt chromatin potential before the affected developmental programs are transcriptionally active, allowing functional abnormalities to emerge only as those programs are deployed later in maturation.

The disproportionate susceptibility of glutamatergic neurons may reflect their developmental state during late gestation, when excitatory projection neurons undergo extensive chromatin remodeling as they transition from migratory to early post-mitotic stages (*53, 54*). This period is characterized by the dynamic reconfiguration of enhancer landscapes that establish lineage-specific transcriptional programs required for circuit assembly. Perturbation of these processes by hypoxia may disrupt enhancer–gene interactions at a critical window, leading to misalignment between regulatory architecture and transcriptional output. Importantly, an acute perturbation of developmental chromatin state need not itself persist unchanged to influence later maturation. A transient disturbance during a critical regulatory transition could alter subsequent developmental trajectories even if the original accessibility state later evolves. Determining which acute chromatin changes persist, resolve, or propagate into later regulatory states will require longitudinal epigenomic profiling across development.

Consistent with these developmental predictions, juvenile animals exposed to prenatal hypoxia exhibited reduced dendritic spine density and increased intrinsic excitability in cortical pyramidal neurons (**Fig. 5, Supplementary Fig. 12**). Similar increased excitability was reported in the entorhinal cortex of rats 3 weeks after transient hypoxia at E14, suggesting that persistent hyperexcitability may generalize across developmental hypoxia paradigms (*55*). In our model, these structural and electrophysiological abnormalities align closely with pathways identified through chromatin accessibility analysis at the fetal stage, rather than with contemporaneous transcriptional changes.

The potassium channel analyses provide a locus-level illustration of this temporal decoupling between early regulatory state and later functional outcome. Although voltage-gated potassium channel genes are minimally expressed in the fetal cortex, their loci exhibit accessible chromatin and are frequently associated with hypoxia-responsive regulatory elements in glutamatergic neurons, consistent with a primed regulatory state (**Fig. 6**). These regions also overlap previously characterized neuronal enhancer elements, supporting their potential regulatory relevance. At later developmental stages, potassium channel-related pathways become transcriptionally dysregulated in the anterior cingulate cortex and are accompanied by reduced sustained potassium currents and altered intrinsic membrane properties. This sequence supports a model in which early perturbations in chromatin accessibility precede and may influence the later deployment of gene-expression programs governing neuronal excitability. Although these data do not establish that individual fetal differentially accessible regions directly cause the later electrophysiological phenotype, they define a coherent sequence in which disruption of a prospective regulatory program precedes dysfunction of the neuronal property it supports.

These findings complement and extend recent work by Riddle et al., who demonstrated that mild neonatal hypoxia without ischemia persistently disrupts synaptic maturation and potassium channel function in the hippocampus (*9*). Whereas that study defined the functional endpoint, persistent synaptic and channel dysfunction after mild hypoxia, the present work identifies an acute upstream regulatory signature at synaptic and ion channel loci that precedes similar functional abnormalities. Together, these studies support a developmental sequence spanning environmental insult, early epigenomic perturbation, and persistent circuit dysfunction. More broadly, our findings suggest that developmental timing fundamentally shapes how environmental injury is encoded. In the developing brain, a regulatory program perturbed by injury may not yet be transcriptionally or functionally active, allowing the consequences of that perturbation to remain phenotypically latent until the program is deployed later in maturation. In this framework, chromatin accessibility functions as a prospective regulatory layer through which transient developmental insults can influence later cellular behavior. Such a mechanism may extend beyond hypoxia to other developmental perturbations, including inflammatory or metabolic stress.

Several limitations shape interpretation and motivate future work. First, we profile a single acute fetal time point; additional fetal and early postnatal time points will be essential to distinguish transient from persistent regulatory changes and to define when (and in which cell types) early chromatin alterations become associated with stable transcriptional consequences. Accordingly, our data do not establish persistence of the fetal accessibility changes themselves; the co-option model instead allows that transient disruption of chromatin state during a critical developmental transition could alter subsequent regulatory trajectories. Second, peak-to-gene linkage and neighboring-gene approaches remain associative; establishing causality will require targeted perturbation of candidate enhancers and cell type-specific manipulation of hypoxia-responsive chromatin regulators. Third, hypoxia was delivered to the pregnant dam, so maternal physiological stress responses may contribute to fetal chromatin changes independently of direct fetal oxygen deprivation; disentangling maternal from fetal-autonomous effects will require targeted approaches. Lastly, low-frequency populations were underpowered for some analyses, and non-cell-autonomous mechanisms, such as immune or vascular signaling, may contribute to later phenotypes in ways not fully captured here.

In summary, transient prenatal hypoxia disproportionately disrupts the chromatin regulatory architecture of developing glutamatergic neurons, altering prospective developmental programs before their functional consequences emerge. These acute chromatin signatures align with later transcriptional, structural, and electrophysiological abnormalities that are not prominent in the immediate transcriptional response, supporting environmental co-option of chromatin potential as a framework linking transient prenatal injury to delayed neurodevelopmental dysfunction. These findings also raise translational questions for the next generation of interventions: whether early chromatin states could eventually inform biomarkers of neurodevelopmental risk and whether modulation of chromatin regulation during or shortly after injury could alter the subsequent trajectory of affected developmental programs.

## MATERIALS AND METHODS

### Animals

Animal care and experimental procedures were performed in accordance with regulations of the Institutional Animal Care and Use Committee of the Children’s Hospital of Philadelphia (protocol IAC 24-000547). All mice were C57BL/6 mice purchased from Charles River Laboratories (Wilmington, MA). Timed matings were used to ensure timing of prenatal hypoxia exposures. The mice were maintained at the Children’s Hospital of Philadelphia Animal Facility at a 12-hour light, 12-hour dark cycle (0615-1815 h), and with ad libitum access to water and diet mouse food 5015 (LabDiet, St. Louis, MO).

### Prenatal Hypoxia

Pregnant dams were exposed to prenatal hypoxia as previously described(*27, 56*). In brief, at embryonic day 17.5 (E17.5) pregnant dams were placed in a controlled oxygen chamber (ProO2110 controller, A-chamber, BioSpherix Ltd., Parish, NY). Mice were acclimated to the chamber for up to 2 minutes at 21% inspired oxygen. For hypoxia, oxygen percentage was decreased with balanced nitrogen to 5% inspired oxygen over 30 minutes and maintained at 5% inspired oxygen for 8 hours. For normoxia, pregnant mice were placed in the same controlled oxygen chamber for 8 hours at 21% inspired oxygen. Dams in both groups were provided free access to food and hydrogel for hydration throughout the time in the chamber. A small tray with sodium lime pellets was utilized to scavenge for exhaled carbon dioxide away from the animals.

### Nuclear Isolation and Sequencing

At the end of exposure, pregnant dams were anesthetized with 5% inhaled isofluorane and terminal hysterectomy was performed. For each fetus, the cortex was individually isolated in PBS on ice and flash frozen and tail was collected for genotyping for sex using Sry. To capture inter-fetal variability while limiting repeated representation within litters, 2–3 cortices from fetuses of the same sex were pooled for each nuclear extraction. Four to five litters were represented per experimental condition, and each litter contributed no more than one male and one female pooled sample. Nuclei from fetal brain were isolated per 10x Genomics protocols. Libraries were prepared using 10x Genomics Chromium Single Cell Multiome ATAC + Gene Expression Reagent Kit per manufacturer’s instructions (Pleasanton, CA). Libraries were uniquely indexed using the Chromium i7 Sample Index Kit, pooled, and sequenced on an Illumina Novaseq 6000 sequencer on an Illumina v1.5_S2 100 cycles flow cell. Libraries were sequenced to a target depth of ∼20,000 paired reads per nucleus for snRNA-seq and ∼25,000 paired reads per nucleus for snATAC-seq. Individual sequencing runs of multi-omic data were analyzed with cellranger-arc-1.0.1 using the mm10 reference genome.

### Computational Analyses

Detailed analyses described below. Packages for R and Python are listed in **Table S4**.

#### Multi-omic Integration

Data were processed using Seurat and Signac for filtering, normalization, dimensional reduction, integration, and analysis(*30, 31, 38, 57*). RNA data were normalized using the standard Seurat workflow, and ATAC data were processed using latent semantic indexing (LSI). Because the ATAC-derived dimensional reduction showed strong segregation by experimental condition, Harmony was applied to the ATAC LSI representation to facilitate joint integration and consistent cell-type annotation across conditions. Harmony-adjusted embeddings were used for multimodal integration, dimensional reduction, visualization, and cell-type annotation only and were not used for differential gene expression or differential accessibility testing. Cell types were annotated through unsupervised clustering and marker gene expression, referencing the Allen Brain Atlas and published datasets(*33, 58*). Major cell classes included glutamatergic neurons, interneurons, neuroprogenitor cells, proliferative cells, astrocytes, oligodendrocyte progenitor cells, microglia, and endothelial cells. Gene activity scores were calculated from snATAC-seq data to infer chromatin accessibility within gene bodies.

#### Single-Nucleus Differential Gene Expression

Wilcoxon rank-sum testing was used to identify differentially expressed genes between hypoxia and normoxia within each annotated cell type using Seurat’s FindMarkers() function (*30, 31, 38, 57*). Individual nuclei were used for cell-type level differential expression testing. Genes with an adjusted p-value < 0.05 and an absolute log2 fold change > 0.25 were considered differentially expressed.

The pathfindR workflow was executed using the run_pathfindR() function with the “GO” gene set database (MSigDb C5 Gene Ontology Collection V 7.4)(*39, 59*). Enrichment results were filtered to retain terms with a FDR < 0.05. We used the cluster_enriched_terms() function to group similar GO terms based on semantic similarity. To quantify the relevance of pathways, we computed a pathway enrichment score using score_terms() for each biological sample for each cell type, calculated the average enrichment score by cell type and used ComplexHeatmap with word cloud to visualize terms(*39, 60*).

Disease- and phenotype-enrichment analyses were performed using DisGeNET through the disgenet2r package, with FDR < 0.05 considered significant (*41*). For visualization, phenotype terms were ranked according to the number of cell-type-specific gene sets in which they reached significance and then by mean FDR across cell types; the top 25 terms were displayed. As a control for gene-set size, random gene lists containing 600–1,000 genes, reflecting the range of DEG-set sizes observed across cell types, were generated and analyzed using the same workflow (generated by https://molbiotools.com/randomgenesetgenerator.php) and also run through disgenet2r. Heatmaps of FDR values were generated using ComplexHeatmap, and phenotype classes were annotated by AGC.

#### Single-Nucleus Dynamical RNA Velocity Estimation

Spliced and unspliced counts were generated using the velocyto “run” command on sorted bam files with reference genome mm10(*61*). Loom files for each sample were generated, and subsequently loom files for each condition were concatenated together to form two looms segregated by condition (i.e., hypoxia exposed, normoxia exposed). We then obtained cell barcodes, expression counts, and dimensionality reduction matrix (UMAP) for both conditions from the Seurat object and converted these data into anndata format in R. Anndata objects were merged with loom files for the appropriate condition, and finally we used scVelo to calculate dynamical RNA velocity for hypoxia and normoxia separately(*62*). Dynamical velocities were plotted on the UMAP coordinates derived from Seurat. Latent time was estimated from dynamical velocities and plotted on UMAP coordinates. Kinetic parameters, including transcription, splicing, and degradation rates, were estimated using the dynamical model.

#### Single-Nucleus Epigenetic Analysis

Fragment files from 10x Genomics cellranger pipeline were imported into ArchR using createArrowFiles()(*45*). To make comparison and naming of nuclei consistent, ArchR object was intersected with metadata from filtered Seurat object. There was a >99% overlap between the two objects. Peak calling was conducted using MACS2 through ArchR’s addReproduciblePeakSet() function, which identifies consensus peaks across grouped cells. Gene scores were computed using addGeneScoreMatrix(), which integrates accessibility signals across gene bodies and promoter regions(*45, 63*).

Differential chromatin accessibility between hypoxia and normoxia was assessed within each annotated cell type using ArchR’s getMarkerFeatures() function, with correction for TSS enrichment and fragment count. Individual nuclei were used for cell-level differential accessibility testing. To focus subsequent analyses on the most robust accessibility changes, differentially accessible regions were defined using stringent thresholds of FDR < 1 × 10−7 and |log2FC| ≥ 1.

Pathway and gene ontology enrichment analyses were performed for genes neighboring differentially accessible regions using Cistrome-GO (http://go.cistrome.org/). Peak-to-gene linkage was performed using ArchR’s addPeak2GeneLinks() function. Peak-to-gene linkage heatmaps were generated using plotPeak2GeneHeatmap() and partitioned into six clusters. Linkage density within each heatmap cluster was quantified by calculating the median density across linked regions for each cell type–condition group. Differences in linkage density between normoxia and hypoxia were assessed within each cell type using permKS() from the R package perm. P values across cell type–cluster comparisons were adjusted with FDR < 0.05 considered significant.

#### Pseudotime Calculation

Pseudotime values were calculated independently for both snRNA and ATAC-seq data using slingshot and condiments(*64, 65*), with neural stem cells specified as the starting population. Pseudotime distributions were compared between normoxia and hypoxia within each lineage using an asymptotic two-sample Kolmogorov–Smirnov test. Calculated pseudotime values were overlaid on the WNN UMAP for visualization.

### Golgi staining

To assess structural changes in cortical pyramidal neurons following prenatal hypoxia, Golgi-Cox staining was performed on brain sections from juvenile mice. Mice were euthanized and brains were processed using the FD Rapid GolgiStain™ Kit (FD NeuroTechnologies, Columbia, MD, USA) according to the manufacturer’s protocol. Briefly, PBS-only perfused brains were maintained in impregnation solution (Solution A and B mixture) for 14 days before transfer to Solution C for 3 days to 1 week in the dark. Brains were then frozen in tissue freezing media and stored in −80°C until sectioning. Coronal sections (150 µm) were cut using a vibratome and mounted on gelatin-coated slides. Layer 5/6 pyramidal neurons in the anterior cingulate cortex were identified at 60x oil immersion magnification in brightfield setting of Keyence microscope. Dendritic spine density was quantified starting approximately 40 µm from the center of the soma, with dendrites tracked in 20-µm increments (median length analyzed = 156.05 µm; range, 34.83–480.92 µm) using Reconstruct (*66*). Spine counts were normalized to dendritic length for each neuron to calculate the spine density. All analyses were performed blind to experimental condition.

### Electrophysiology Cortical Slice Preparation

Acute cortical slices were prepared as previously described(*67*). Briefly, postnatal day (PND) 25-30 male or female mice previously exposed to normal air (normoxia) or air with 5% inspired oxygen (hypoxia) for 8 hours in utero at embryonic day 17.5 were anesthetized with isoflurane, transcardially perfused with iced cold (0–4 °C) sucrose artificial cerebrospinal fluid (aCSF) containing in (mM): sucrose 204, KCl 2.5, CaCl_2_ (Anhydrous) 0.5, MgSO_4_ (Anhydrous) 10, NaH_2_PO_4_ 1.25, NaHCO_3_ 26 and glucose 11, Na-Ascorbate 5, Na-Pyruvate 3, and Thiourea 2. The whole brain was rapidly removed and immersed in ice-slush sucrose aCSF for 2-3 min. Coronal slices (350µm) of the cortex were prepared in the same ice-slushy sucrose aCSF using a Leica vibratome (VT1200, Leica, Germany). Before recordings, slices were allowed to recover at 32°C in recovery aCSF (115 mM NaCl, 2.5 mM KCl, 1mM MgSO_4_, 2.5 mM CaCl_2_, 1.4 mM NaH_2_PO_4_, 30.8 mM NaHCO_3_, 15.5 mM glucose, 5mM NaAscorbate, 3mM NaPyruvate, and 2mM Thiourea) for at least 1h prior to whole-cell recording. Following recovery, slices were transferred to a submerged chamber (RC-27, Warner Instruments) and continuously perfused with oxygenated recording aCSF (128mM NaCl, 2.5 mM KCl, 1mM MgSO_4_, 2.5mM CaCl_2_, 1.4mM NaH_2_PO_4_, 30mM NaHCO_3_, 15.5mM glucose) with a flow rate of 2 mL/min using a gravity-driven perfusion system. All solutions were equilibrated with 95% O_2_/5% CO_2_ to maintain a pH near 7.4 and all recordings were performed at 32°C.

### Electrophysiological Recording

Following recovery, cortical slices were transferred to a chamber on a BX-51WI microscope (Olympus, USA) equipped with IR-DIC and fluorescent optics, an IR-sensitive CCD camera (Rolera, Qimaging), and a 40× water immersion objective. Slices were visualized on a screen and whole-cell recordings were made on pyramidal cells (PCs) in the anterior cingulate cortex (ACC). PCs were targeted based on their pyramidal-like shape and clear apical dendrite. Patch pipettes (4–6 ΜΩ) were pulled from thick-walled borosilicate glass (ID: 1.1 mm, OD: 1.5; WPI, Sarasota, FL) using a horizontal puller (Sutter Instruments P-97) and filled with (in mM) 5 KCl, 135 K-gluconate, 2 NaCl, 10 HEPES, 4 EGTA, 4 MgATP, and 0.3 Na_3_GTP (pH ≈ 7.2; 285 mOsm). The liquid junction potential between the recording aCSF and the internal solution was cancelled out using the bridge-balancing circuit of the amplifier before seal formation. All recordings were performed with a Multiclamp 700B amplifier (Molecular Devices, USA) connected to a Digidata 1440A analog-digital converter (Molecular Devices, USA) and signals were sampled at 20 kHz and filtered with a 5 kHz low-pass Bessel filter. To reduce the impact of artifacts on the recordings, series resistance and capacitance compensation (70–80%) were performed online to decrease the transient observed under a rectangular test pulse. Patch quality was monitored throughout the experiments and only recordings with stable holding current (<100 pA) and access resistance (<30 MΩ) were used for analysis. Upon establishing whole-cell configuration on pyramidal cells, we measured the membrane response to subthreshold current injection in current-clamp mode using steps ranging from 20 to −100 pA, with a duration of 250 ms and a frequency of 10Hz. The −20 pA step was used to calculate the input resistance because this current intensity did not produce membrane voltage sag. Action potential firing properties were subsequently characterized in response to a 100 or 200pA step current of durations ranging from 50-1000 ms(*68*). To record voltage-gated sodium (Na^+^) and potassium (K^+^) currents, a conventional step protocol (500ms step duration from −90 to +50 mV in 10 mV increments at 5 Hz) was used in the voltage-clamp mode. The fast-activated deflections at the beginning of these voltage steps were attributed to transient sodium channel currents, and the quasi-steady-state currents at the end of the voltage steps were attributed to potassium channel currents (*69*). All data were analyzed using Clampfit 10.7 (Molecular Devices, USA) or Minianalysis 6.0.3 (Jaejin software, Leonia, NJ, USA).

### Bulk RNA-sequencing and Analysis

Twelve juvenile brains (P27-P28) evenly split between males and females that represented 6 litters were microdissected for the cingulate cortex. Sequencing was performed at the CHOP High Throughput Sequencing Core. Briefly, RNA was isolated using the QIAGEN RNEasy kit (Hilden, Germany), with 260/280 ratios indicating high RNA quality (range 2.06-2.11). About 500 ng was sequenced on an Illumina NextSeq 2000 using poly (A)- enriched mRNA, with paired-end 150 bp reads, yielding approximately 30-50 million reads per sample. Fastq files were processed using BioJupies (v1.1.6) to the mm10 reference genome for EnrichR gene ontology analysis (*70, 71*).

### Statistical analysis

Statistical analyses were performed using Excel and GraphPad Prism software (San Diego, CA, USA). Nested t-test was used to calculate statistical significance for dendritic spines. Statistical significance for action potential number relative to step duration and I-V curves were determined by the Two-way ANOVA test with a Šídák’s multiple comparisons post hoc test. The Student *t*-test was used to assess statistical significance for input resistance analysis. Data are given as means (± SEM) and differences in means were considered statistically significant at p˂0.05. Significance levels [p<0.05 (*), p<0.01(**), p<0.001(***), p<0.0001(****)] are indicated in the text and figures. Sample sizes were not predetermined by power analysis, but they conformed to those reported in similar studies. A chi-square test was used to compare the proportion of potassium channel genes with neighboring differentially accessible regions with that of randomly selected genes.

## Acknowledgements

We thank the CHOP Center for Applied Genomics and High Throughput Sequencing Core for sample processing and sequencing for single–nucleus multi–omic data and bulk RNA–seq, respectively. Single–nucleus multi-ome sequencing was funded by the CHOP K–readiness Pilot Award. We thank Ima Samba for pilot data for the Golgi staining protocol and Dr. Michal Elovitz, Dr. Mariko Bennett, and members of the Cristancho lab for discussions and thoughtful review of the manuscript.

## Author contributions

Conceptualization: A.G.C.

Methodology: A.G.C., D.J.J., M.M.C., P.S.C., E.J.G., E.C.G., D.Z., B.N.R., E.D.M.

Investigation: A.G.C., D.J.J., M.M.C., P.S.C., E.J.G., E.C.G., D.Z., B.N.R.

Visualization: A.G.C., D.J.J., M.M.C.

Supervision: A.G.C., E.D.M.

Writing—original draft: A.G.C

Writing—review & editing: A.G.C., D.J.J., M.M.C., P.S.C., E.J.G., E.C.G., D.Z., B.N.R., E.D.M.

## Data, code, and materials availability

Data will be deposited in GEO and made publicly available upon publication; accession numbers will be provided. All novel code will be provided upon request. No new packages were developed for this analysis.

## Supplementary Materials

Supplementary material is available online and includes additional figures and tables.

