## Supplemental Figures and Tables for "Environmental co-option of chromatin potential links prenatal hypoxia to persistent cortical circuit dysfunction"

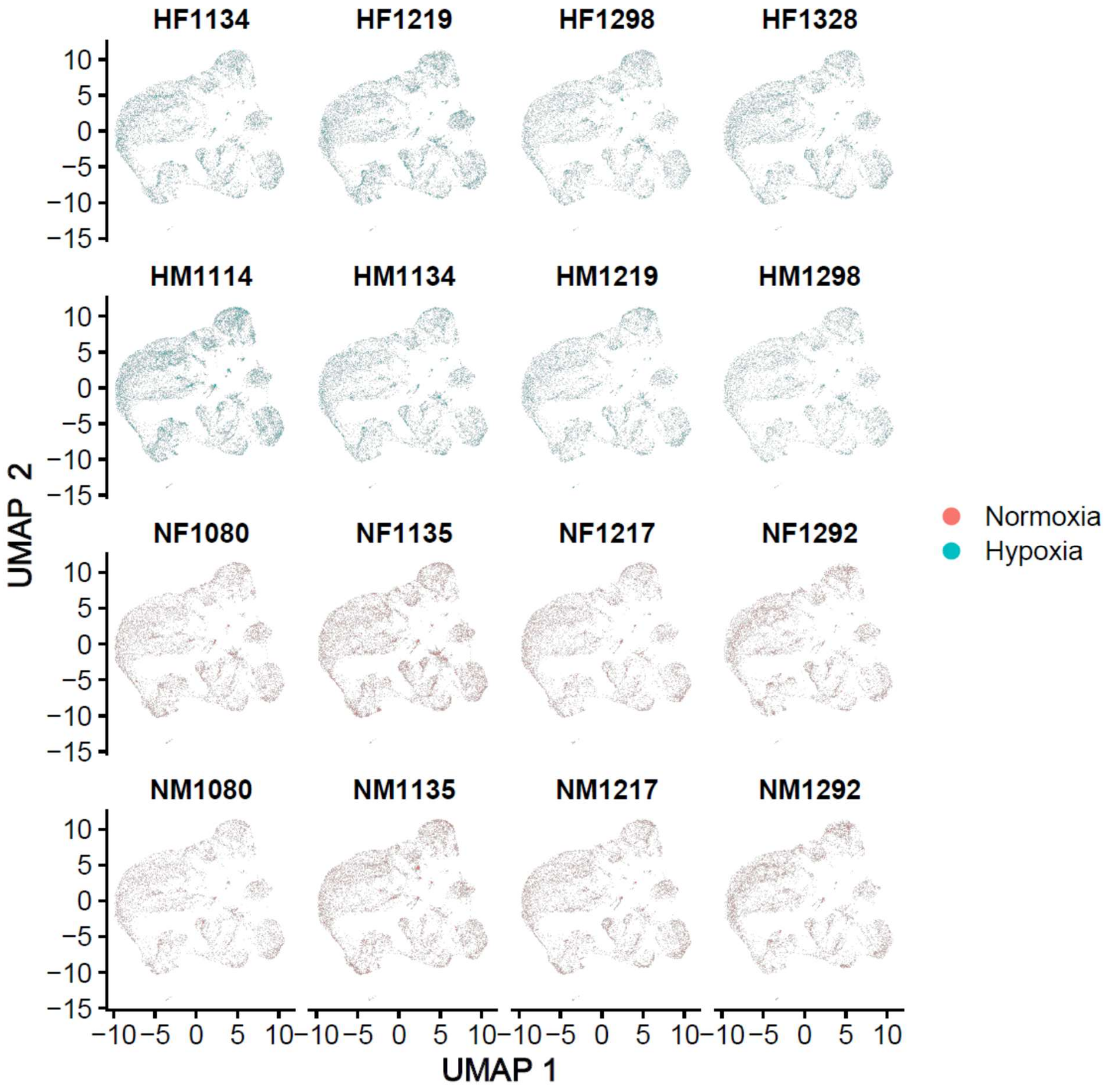

**Fig. S1.Sample-level snRNA-seq dimensionality reduction before multimodal integration.** UMAP projections of snRNA-seq data shown separately for each of the 16 biological samples. Samples are labeled by condition and sex (H, hypoxia; N, normoxia; F, female; M, male). Points represent individual nuclei. Hypoxia samples are shown in turquoise and normoxia samples in coral.

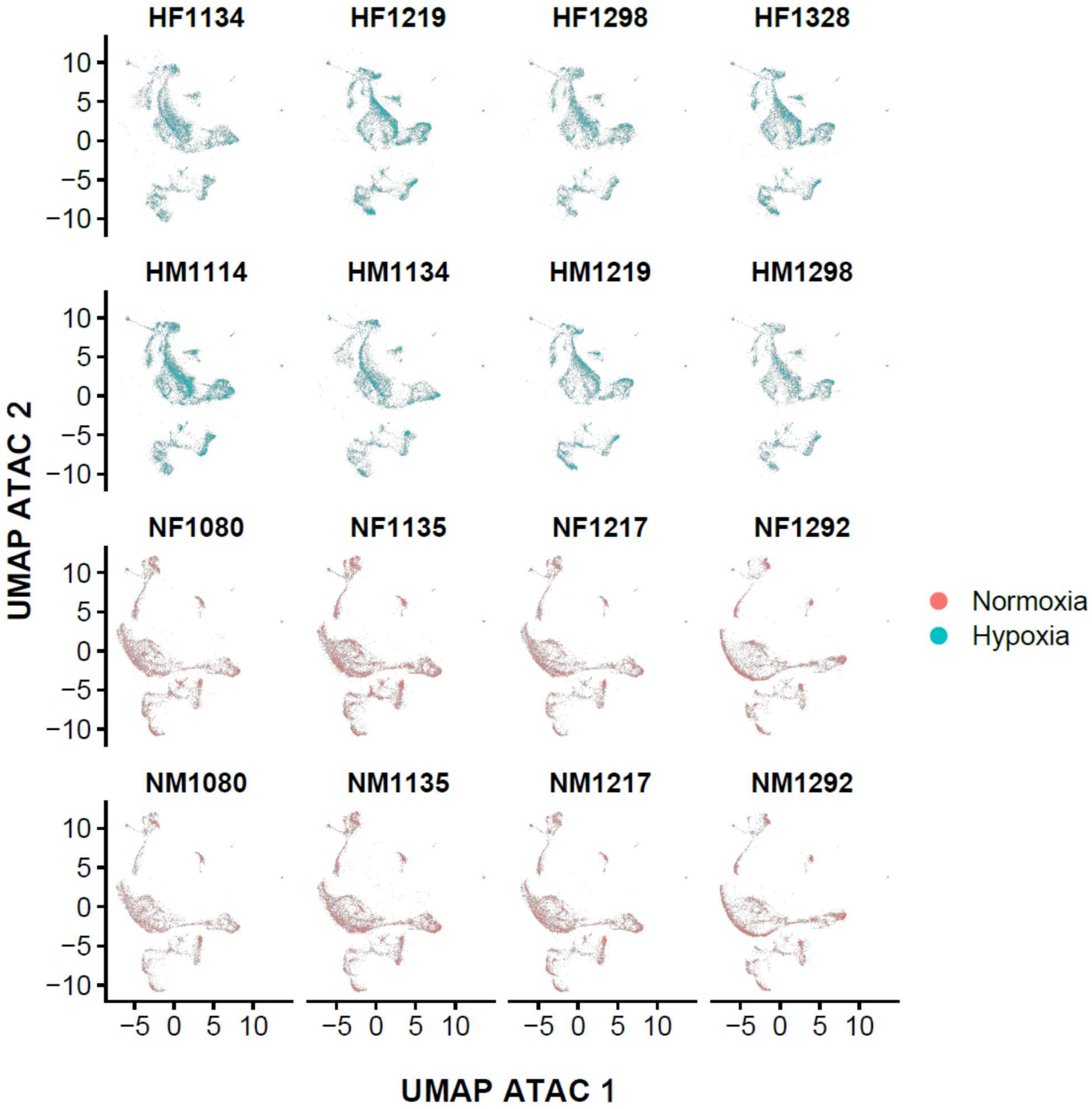

**Fig. S2. Sample-level snATAC-seq dimensionality reduction before Harmony integration.** UMAP projections derived from snATAC-seq data shown separately for each of the 16 biological samples. Samples are labeled by condition and sex (H, hypoxia; N, normoxia; F, female; M, male). Points represent individual nuclei. Hypoxia samples are shown in turquoise and normoxia samples in coral.

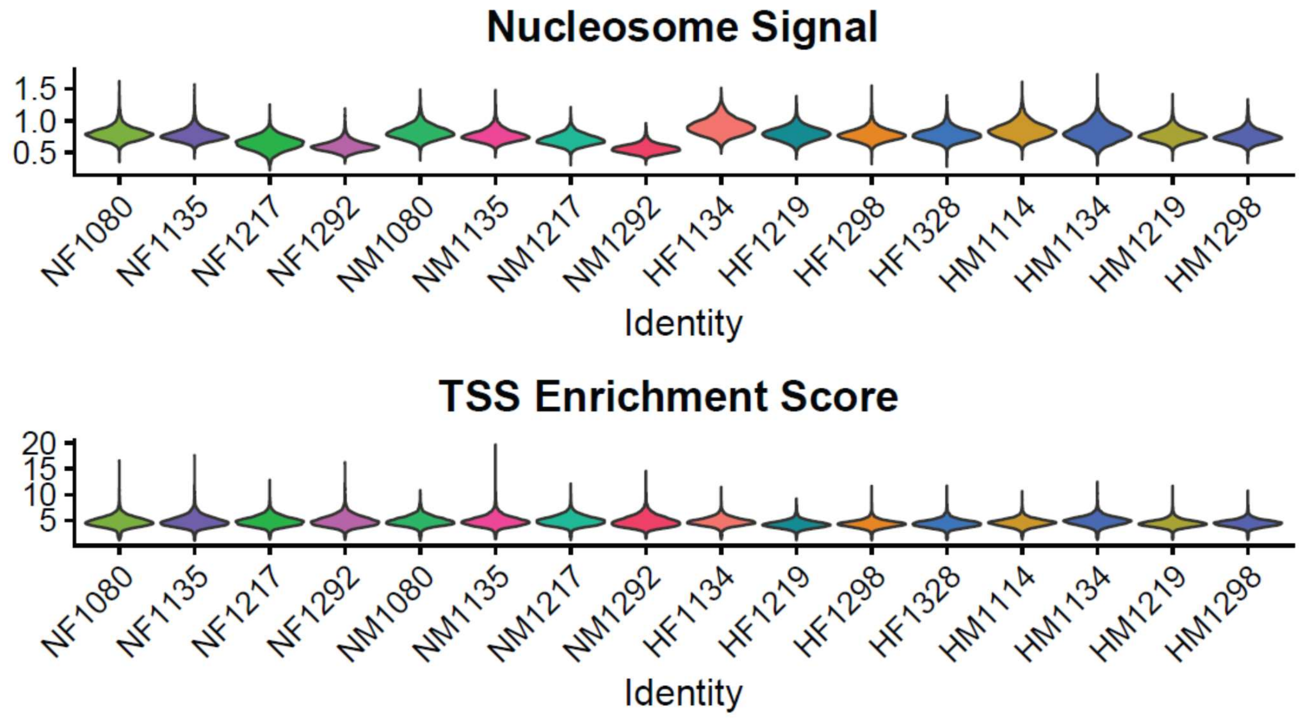

**Fig. S3: snATAC-seq quality-control metrics across biological samples.** Violin plots showing nucleosome signal (top) and transcription start site (TSS) enrichment score (bottom) for each sample. Distributions were comparable across experimental groups, arguing against global differences in ATAC-seq quality as the primary source of condition-dependent separation.

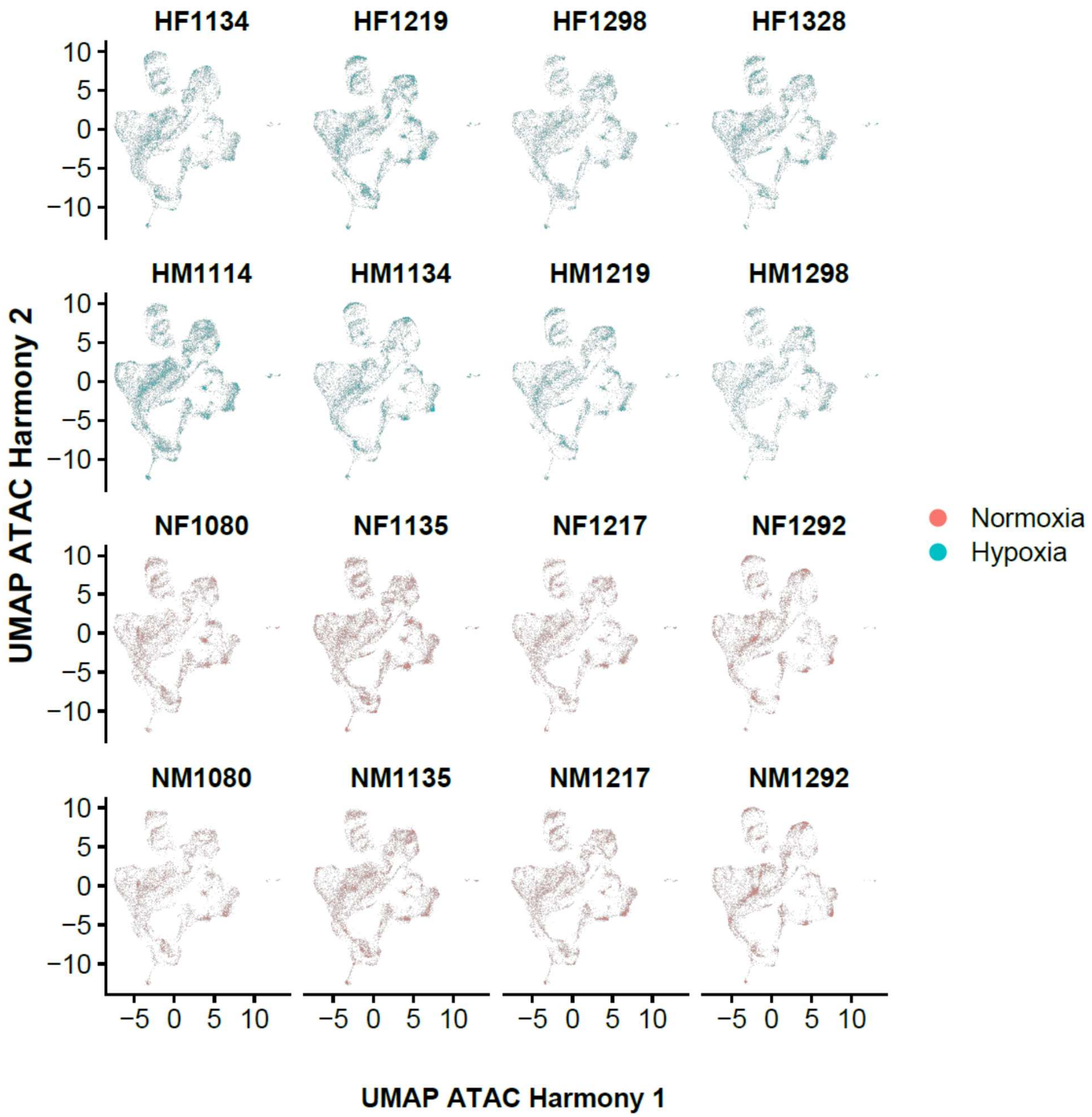

**Fig. S4. Sample-level snATAC-seq dimensionality reduction after Harmony integration.** Harmony-adjusted ATAC UMAP projections shown separately for each biological sample. Samples are labeled by condition and sex as in Fig. S2. Harmony-adjusted embeddings were used for multimodal integration, visualization, and cell-type annotation but not for differential accessibility analyses.

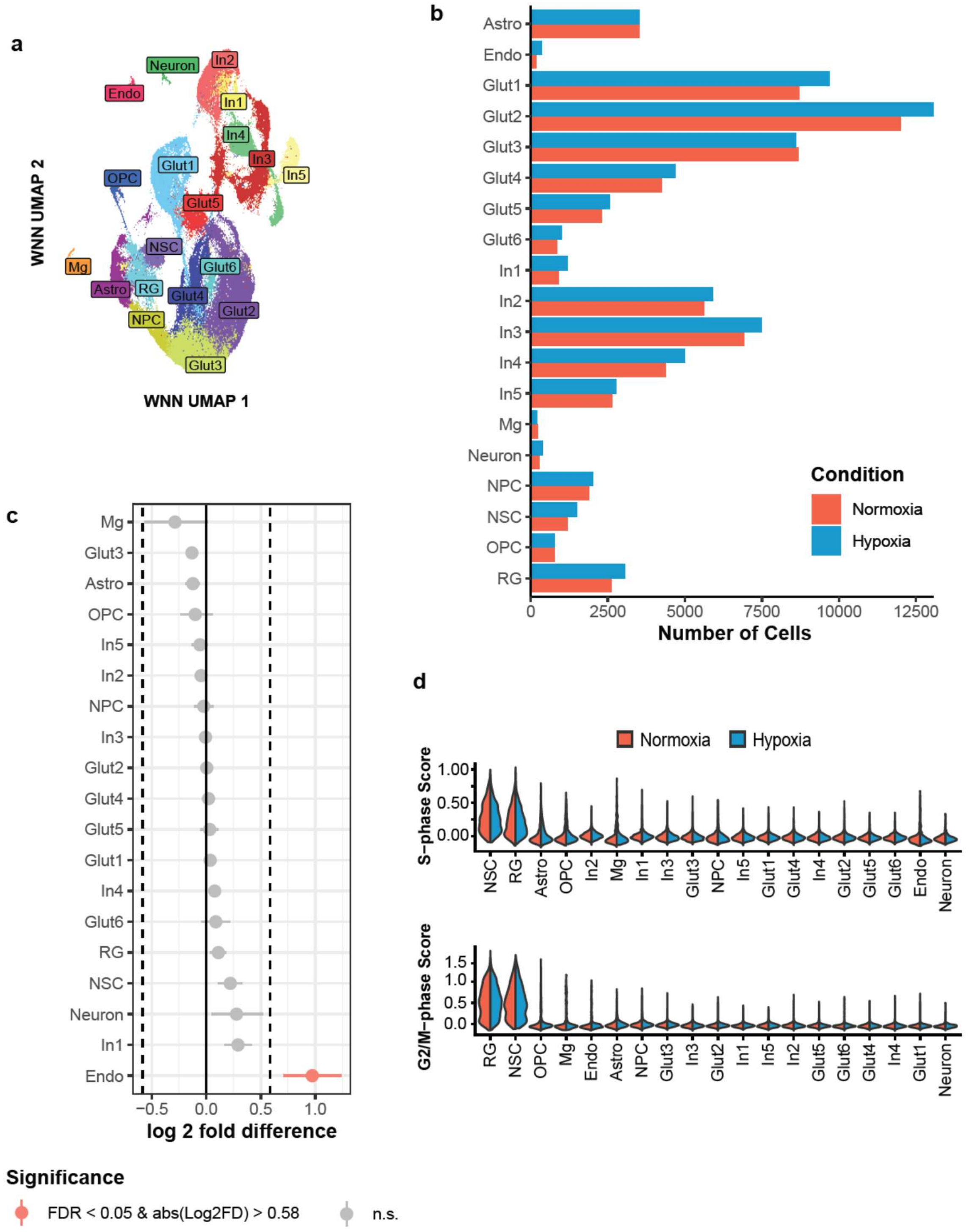

**Fig. S5: Cell composition and cell-cycle characteristics of the integrated snRNA/snATAC-seq dataset. (A)** WNN UMAP of the integrated fetal cortical dataset labeled by annotated cell population. **(B)** Number of nuclei assigned to each cell type in normoxia and hypoxia. **(C)** Differences in cell-type representation between conditions assessed by permutation testing. Points indicate the log2 fold difference between hypoxia and normoxia; significant differences are shown in red and nonsignificant differences in gray. **(D)** S-phase and G2/M-phase cell-cycle scores across cell types, shown separately for normoxia and hypoxia.

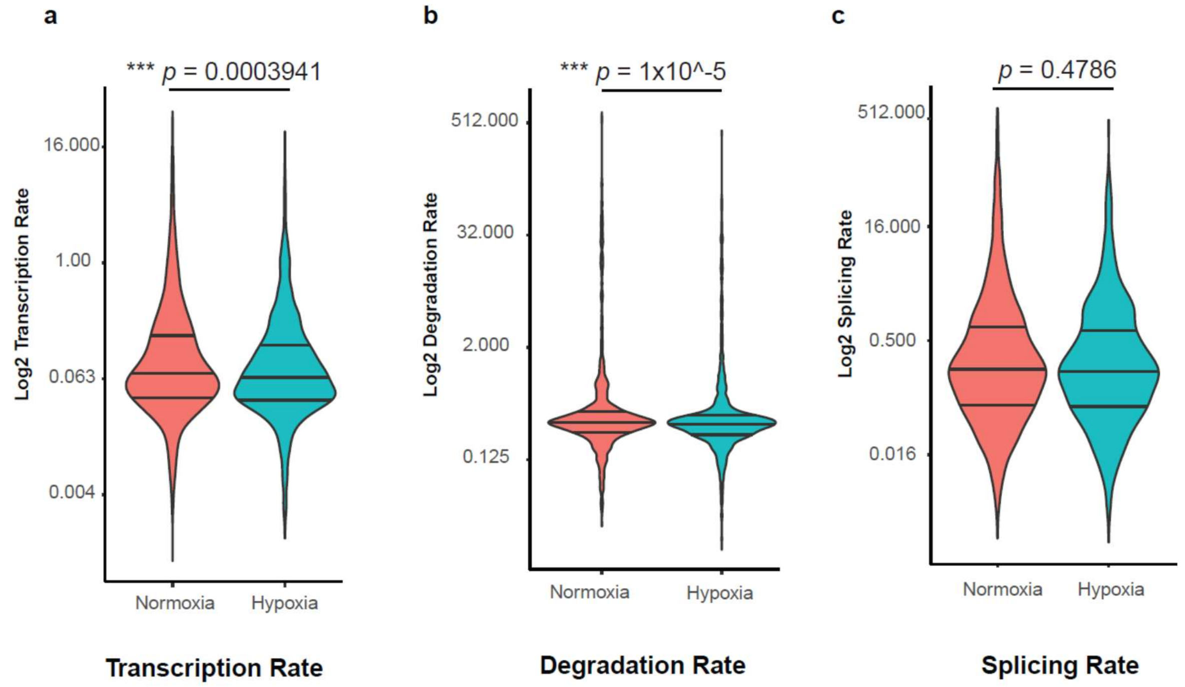

**Fig. S6: Hypoxia alters RNA kinetic parameters.** Violin plots of gene-wise kinetic parameters estimated from the dynamical RNA-velocity model in normoxia and hypoxia: **(A)** transcription rate, **(B)** degradation rate, and **(C)** splicing rate. Transcription and degradation rates were reduced following hypoxia, whereas splicing rates did not differ significantly between conditions. Statistical comparisons were performed using the Wilcoxon test; transcription, $p = 0.0003941$; degradation, $p = 1 \times 10^{-5}$; splicing, $p = 0.4786$.

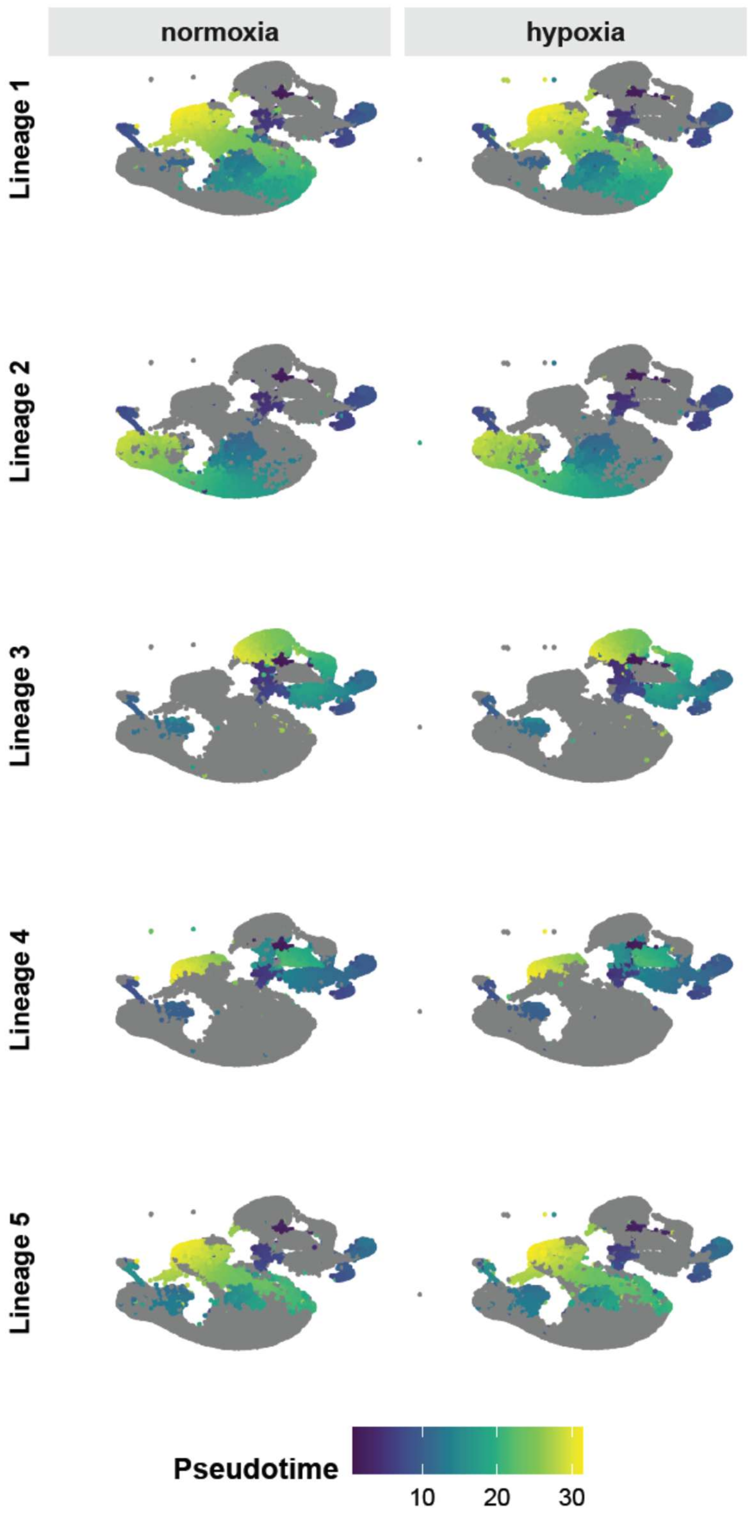

**Fig. S7: RNA-based developmental trajectories are largely preserved following prenatal hypoxia.** snRNA-seq pseudotime projected onto the WNN UMAP for five independently inferred cortical lineages originating from the neural stem cell population, shown separately for normoxia and hypoxia. Cells belonging to each inferred lineage are colored by pseudotime; nuclei outside the indicated trajectory are shown in gray.

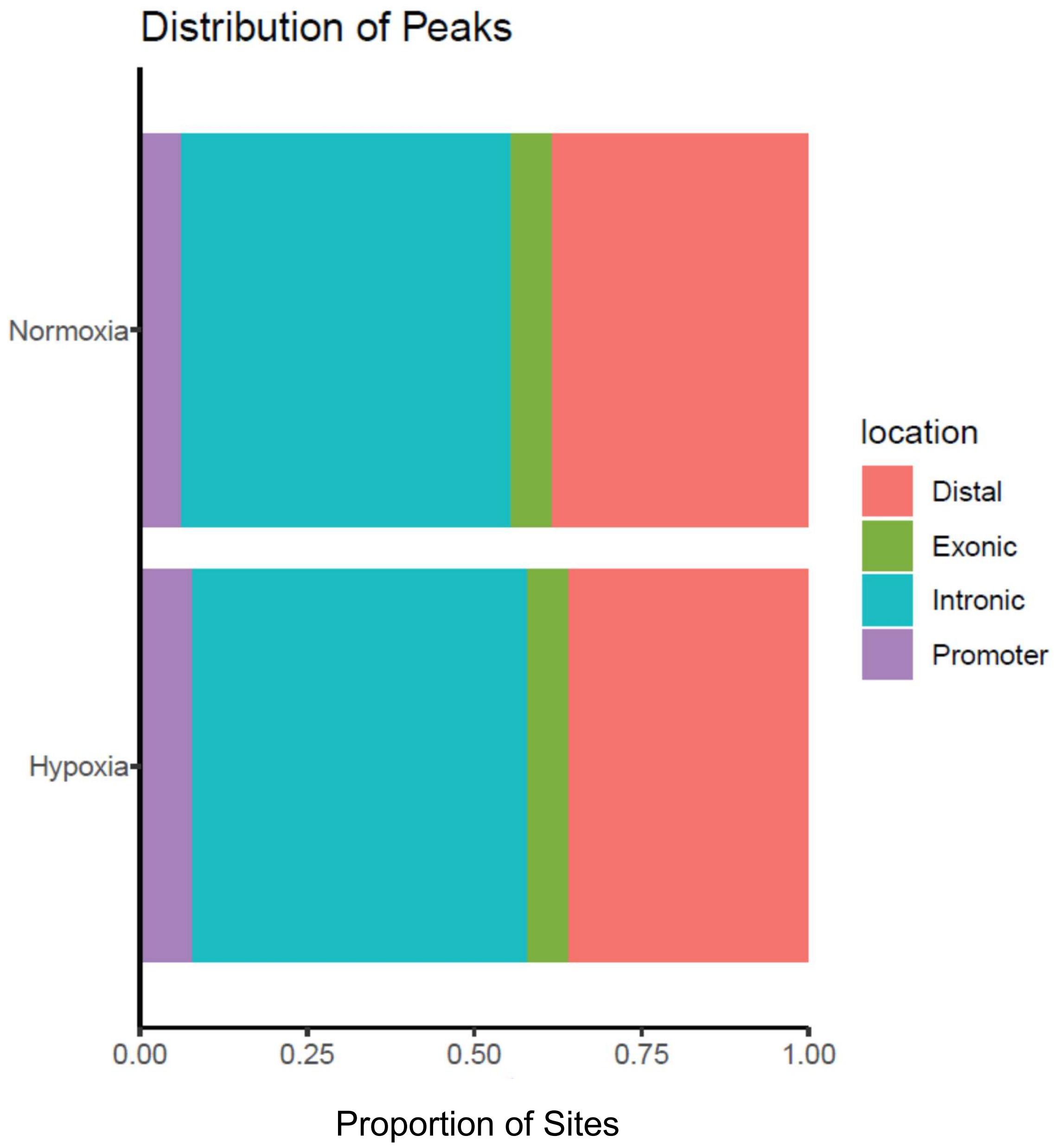

**Fig. S8: Genomic distribution of accessible chromatin regions in normoxia and hypoxia.** Proportion of detected snATAC-seq peaks classified as promoter, intronic, exonic, or distal regions in each condition. Promoters were defined as 2,000 bp upstream to 100 bp downstream of the transcription start site.

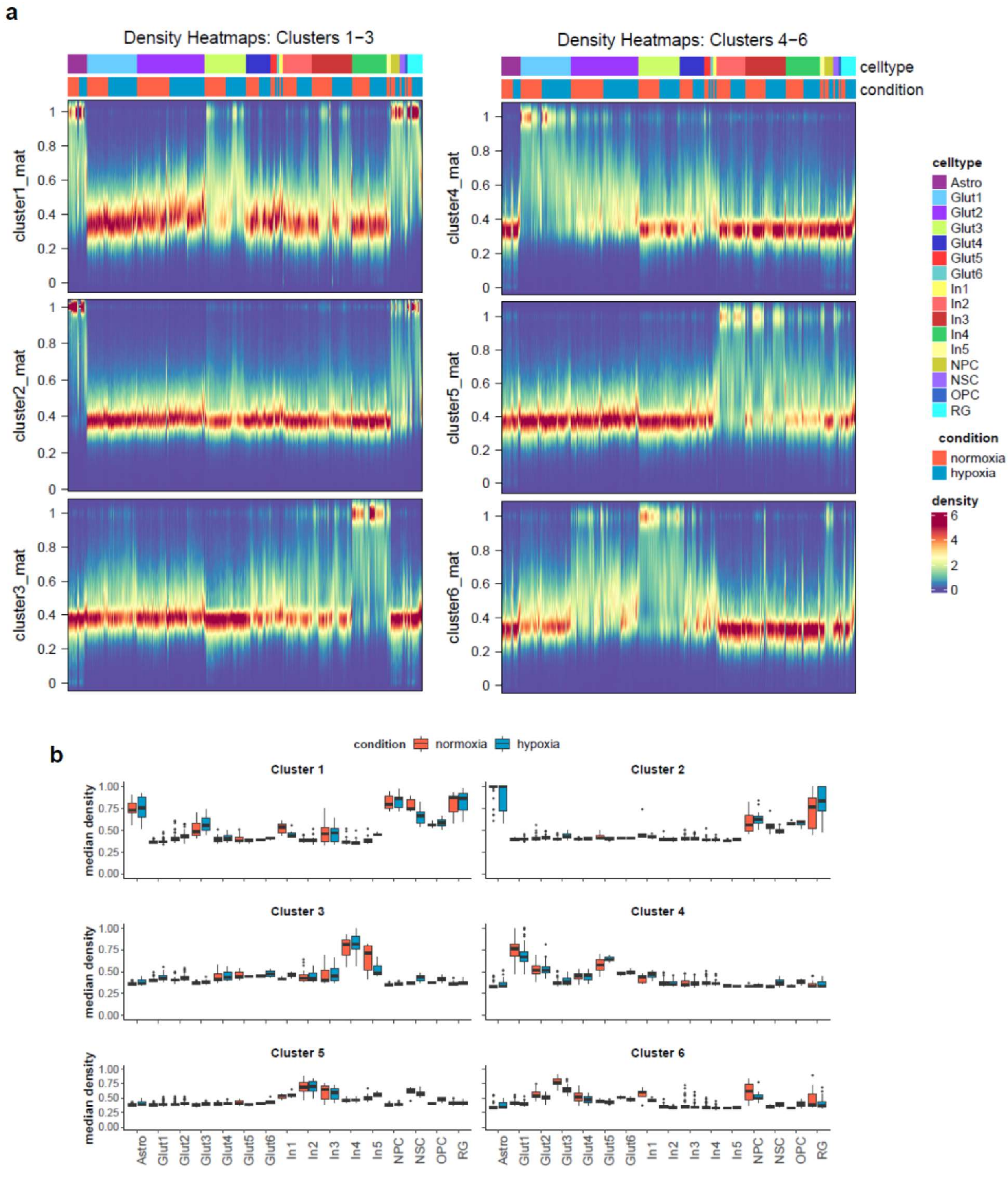

**Fig. S9: Quantification of condition-dependent differences in peak-to-gene linkage density.** **(A)** Density representations of the six clusters identified in the peak-to-gene linkage heatmap, with columns annotated by cell type and condition. **(B)** Median linkage density within each cluster, stratified by cell type and experimental condition. Box plots compare normoxia and hypoxia for each cell type and were used for the permutation-based comparisons summarized in Fig. 4C.

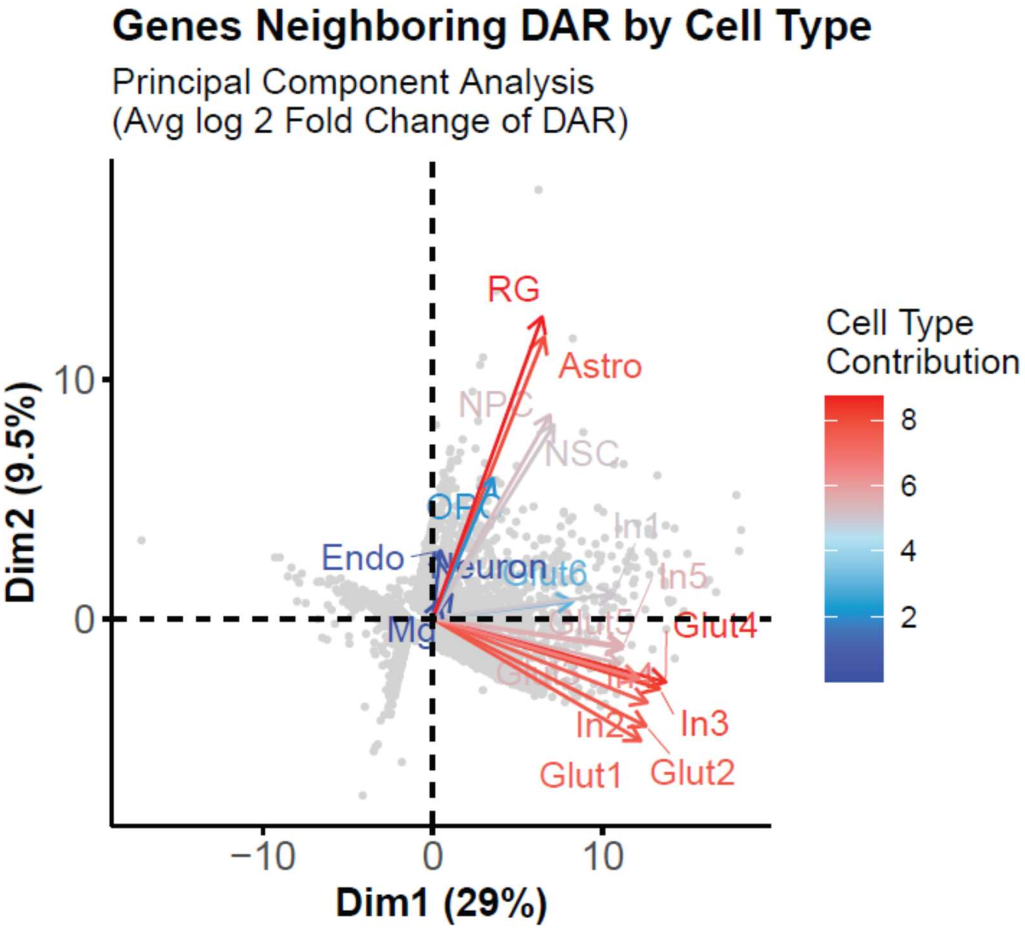

**Fig. S10: Fig. S10. Cell type-specific patterns of differential chromatin accessibility.** Principal component analysis of genes neighboring hypoxia-associated differentially accessible regions across cortical cell types, based on the average log2 fold change in accessibility of neighboring regions. Vectors indicate the contribution of each cell type to the principal-component space.

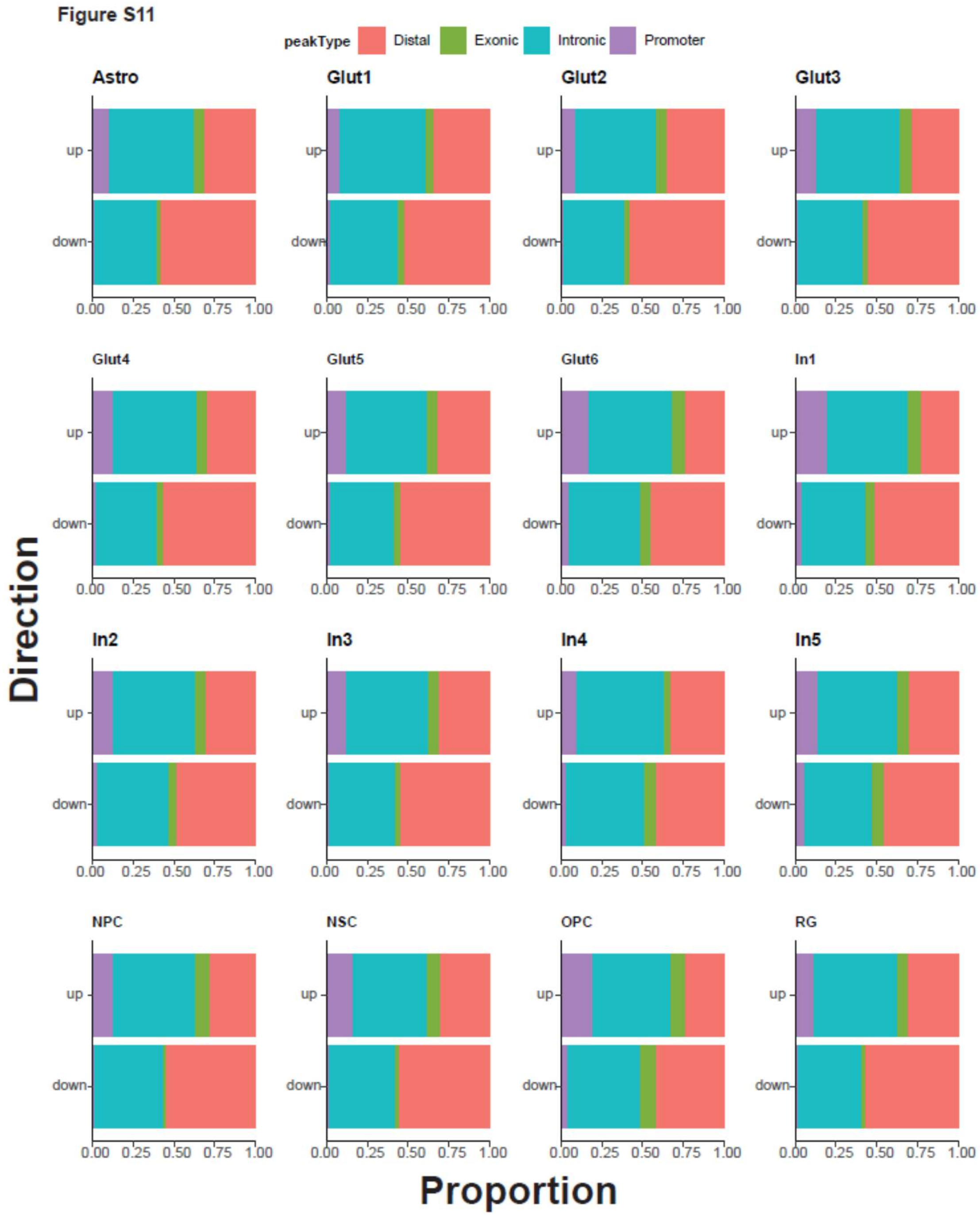

**Fig. S11: Genomic distribution of hypoxia-responsive differentially accessible regions across cell types.** Proportional distribution of differentially accessible regions classified as promoter, intronic, exonic, or distal for each cortical cell population. Regions with increased accessibility following hypoxia are shown as "up," and regions with decreased accessibility as "down."

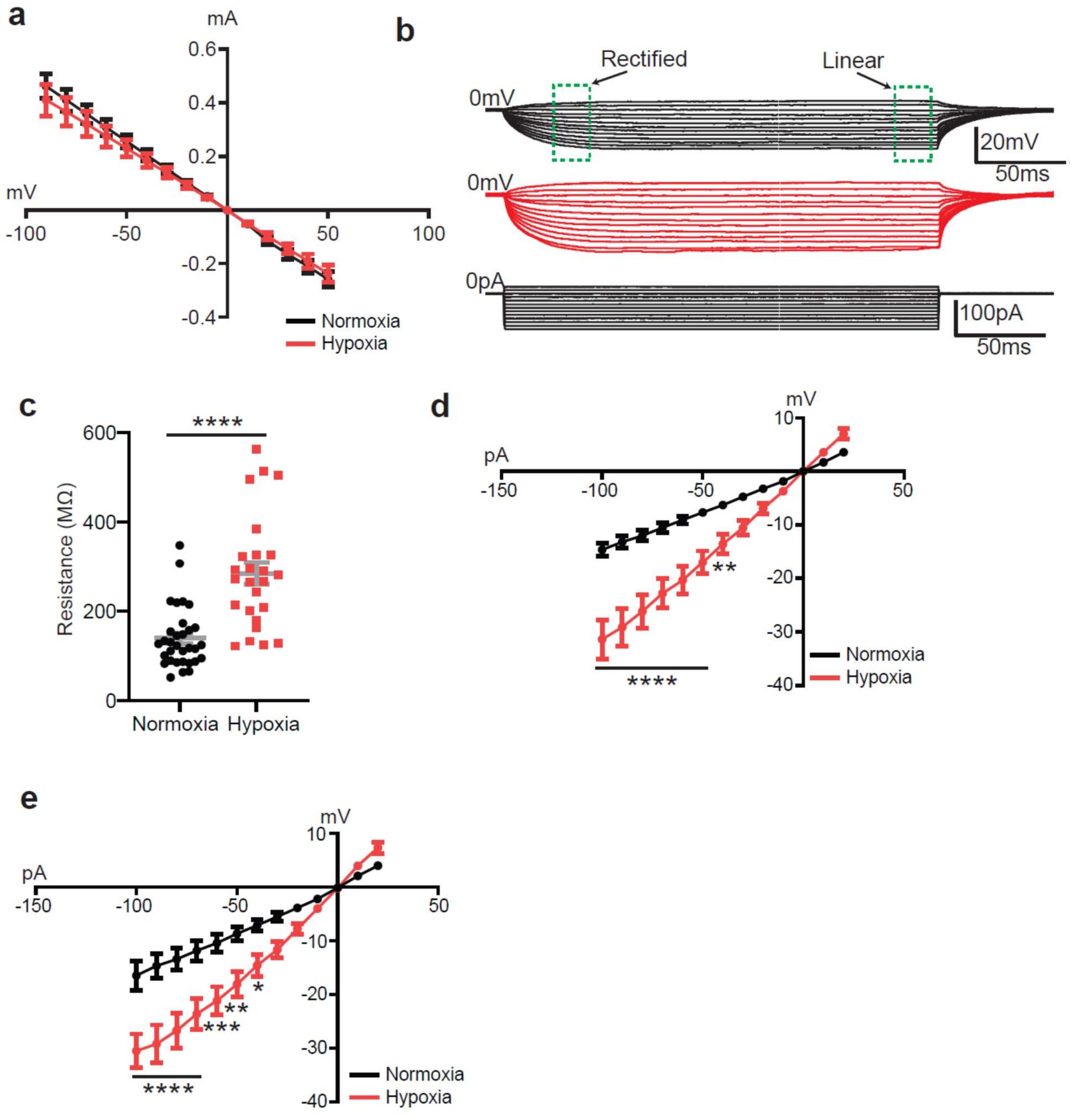

**Fig. S12**. **Prenatal hypoxia alters passive membrane properties of juvenile cortical pyramidal neurons. (A)** Peak current-voltage (I–V) relationship in normoxia and hypoxia [n = clarify recording units/animals]; no significant condition effect was detected [$F(1,715) = 1.048$, $p = 0.3062$]. **(B)** Representative current-clamp traces evoked by subthreshold current steps, with rectified and linear ranges indicated. **(C)** Input resistance was increased following prenatal hypoxia (normoxia, 139.2 ± 12.0 MΩ; hypoxia, 284.4 ± 25.3 MΩ; $p < 0.0001$). **(D,E)** I–V relationships for subthreshold membrane deflections in the rectified (D) and linear (E) ranges. Data are mean ± SEM. I–V curves were analyzed by two-way ANOVA followed by Šídák multiple-comparisons testing. *$p < 0.05$, **$p < 0.01$, ***$p < 0.001$, ****$p < 0.0001$.

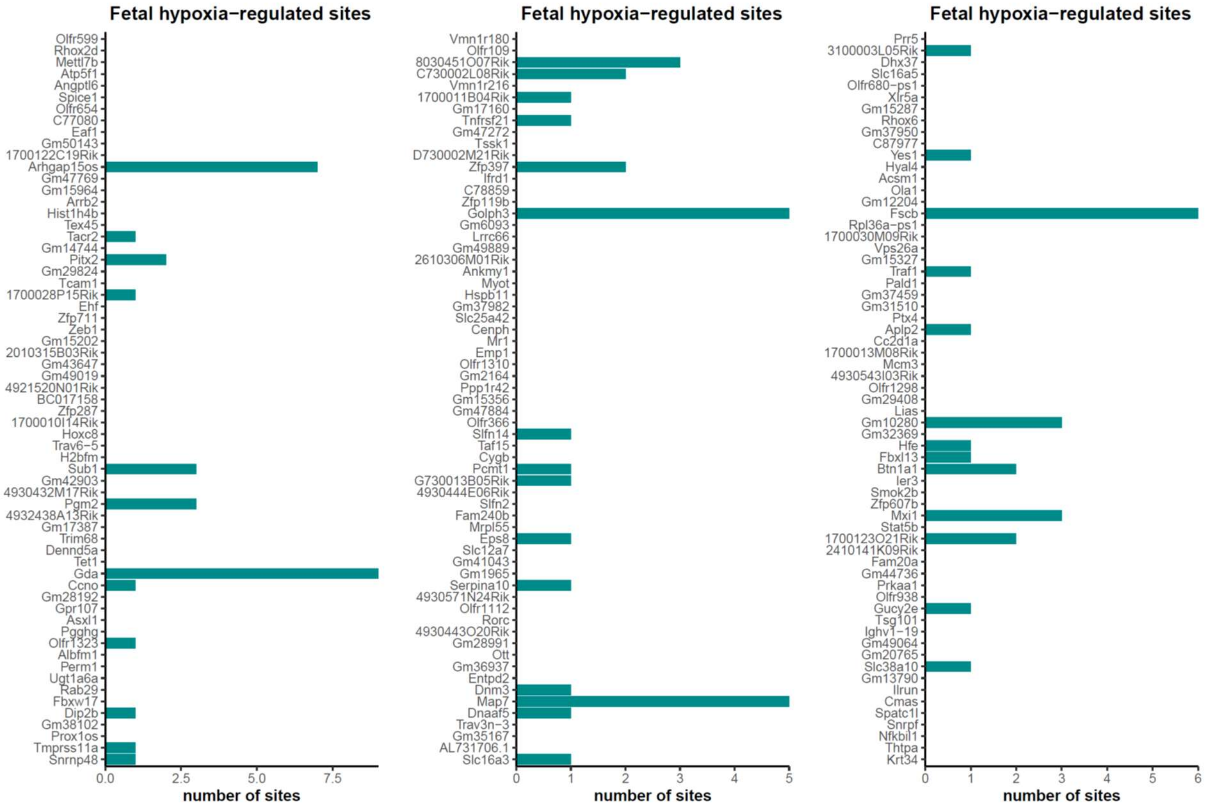

**Fig. S13: Hypoxia-responsive chromatin regions neighboring randomly selected genes.** Three size-matched independent sets of 63 genes expressed in the fetal cortical dataset were randomly selected for comparison with the 63 potassium channel genes analyzed in Fig. 6. Bars indicate the number of neighboring hypoxia-responsive differentially accessible regions associated with each randomly selected gene. These random gene sets were used to assess whether potassium channel genes were disproportionately associated with hypoxia-responsive chromatin regions.

**Table S1: Sample Information for sequencing**

| Sample name | Alt name | Dam | O2 condition | Sex | Nuclei concentration loaded | Number of nuclei loaded | Number of sequenced cells | Median high quality ATAC Fragments per nucleus | GEX median genes per nucleus | Filtered cell number |
|---|---|---|---|---|---|---|---|---|---|---|
| 1_1292_F_Multiome | NF1292 | 1292 | Normoxia | Female | 6,670 | 16,008 | 8,939 | 8,865 | 2,408 | 8,129 |
| 1135_F_Multiome | NF1135 | 1135 | Normoxia | Female | 4,835 | 16,100 | 13,321 | 6,823 | 2,096 | 11,381 |
| 1080_F_Multiome | NF1080 | 1080 | Normoxia | Female | 7,505 | 15,761 | 10,207 | 7,680 | 2,125 | 8,642 |
| 1217_F_Multiome | NF1217 | 1217 | Normoxia | Female | 3,505 | 16,123 | 9,119 | 7,136 | 2,373 | 7,699 |
| 1292_M_Multiome | NM1292 | 1292 | Normoxia | Male | 6,130 | 15,938 | 9,217 | 8,485 | 2,470 | 8,271 |
| 1217_M_Multiome | NM1217 | 1217 | Normoxia | Male | 7,495 | 15,740 | 9,572 | 8,639 | 2,190 | 8,412 |
| 1135_M_Multiome | NM1135 | 1135 | Normoxia | Male | 6,345 | 16,100 | 11,680 | 7,647 | 2,316 | 9,871 |
| 1080_M_Multiome | NM1080 | 1080 | Normoxia | Male | 7,565 | 15,887 | 7,351 | 10,275 | 2,304 | 6,581 |
| 1134_M_Multiome | HM1134 | 1134 | Hypoxia | Male | 6,890 | 15,847 | 9,835 | 9,872 | 2,282 | 8,958 |
| 1298_M_Multiome | HM1298 | 1298 | Hypoxia | Male | 5,260 | 16,306 | 7,200 | 11,004 | 2,254 | 6,588 |
| 1114_M_Multiome | HM1114 | 1114 | Hypoxia | Male | 8,590 | 16,100 | 15,800 | 5,445 | 1,658 | 13,631 |
| 1219_M_Multiome | HM1219 | 1219 | Hypoxia | Male | 7,735 | 16,244 | 8,655 | 9,949 | 2,201 | 7,923 |
| 1134_F_Multiome | HF1134 | 1134 | Hypoxia | Female | 9,405 | 15,989 | 10,303 | 8,913 | 2,203 | 9,470 |
| 1298_F_Multiome | HF1298 | 1298 | Hypoxia | Female | 8,765 | 15,777 | 9,332 | 9,750 | 2,028 | 8,403 |
| 1328_F_Multiome | HF1328 | 1328 | Hypoxia | Female | 9,135 | 16,443 | 9,944 | 9,546 | 2,190 | 9,051 |
| 1219_F_Multiome | HF1219 | 1219 | Hypoxia | Female | 3,475 | 14,595 | 12,765 | 8,084 | 2,104 | 11,373 |
| | | | | **Median** | 7,192.50 | 15,998.50 | 9,703.50 | 8,752.00 | 2,202.00 | 8,527.00 |
| | | | | **St Dev** | 1,835.82 | 409.50 | 2,227.95 | 1,453.33 | 187.83 | 1,832.37 |

**Table S2: Cluster Annotations by RNA-seq Expression Markers**

| Cluster | Class | Top 10 Subclass Markers | Other Subclass Markers | Annotation | Expanded Cell types |
|---|---|---|---|---|---|
| 1 | Neurod2, Rbfox1 | Unc5d, Sema3c | Satb2, Slc17a6 | Glut3 | Glutamatergic 3 (Migrating Pyramidal Neurons 1) |
| 2 | Neurod2, Neurod6, Rbfox1 | Tle4, Zfpm2 | Ldb2 | Glut1 | Glutamatergic 1 (Corticothalamic Neurons) |
| 3 | Gad2 | Sst, Nrxn3 | Lhx6, Sema3a, Reln | In2 | Interneurons 2 (Lhx6/Sst+) |
| 4 | Neurod2, Neurod6, Rbfox1 | Satb2 | Dok5, Ldb2 | Glut2 | Glutamatergic 2 (Callosal Projection Neurons) |
| 5 | Neurod6, Rbfox1 | Satb2 | Dok5, Nrxn3, Ldb2 | Glut2 | Glutamatergic 2 (Callosal Projection Neurons) |
| 6 | Rbfox1, Rbfox3 | | Zfpm2, Sema3a, Slc17a6 | Glut4 | Glutamatergic 4 (Migrating Pyramidal Neurons 2) |
| 7 | Gad2 | | Isl1, Ldb2 | In4 | Interneurons 4 (Isl1+) |
| 8 | Gad2 | Adarb2, Nrxn3, Dlx6os1 | | In3 | Interneuron 3 (Adarb2+) |
| 9 | | Slc1a3, Fabp7 | Zfpm2, Nrxn3 | Astro | Astrocytes |
| 10 | Gad2 | | Drd2, Tle4 | In5 | Interneurons (Drd2+) |
| 11 | Neurod6, Rbfox1 | Satb2 | Dok5, Ldb2 | Glut2 | Glutamatergic 2 (Callosal Projection Neurons) |
| 12 | Rbfox1 | Ldb2 | Dok5 | Glut5 | Glutamatergic 5 (Unspecified Excitatory Neurons 1) |
| 13 | Neurod2, Neurod6 | Unc5d | Sema3c, Satb2, Sema3a, Slc17a6 | Glut3 | Glutamatergic 3 (Migrating Pyramidal Neurons 1) |
| 14 | Gad2 | | Adarb2, Sema3a, Slc17a6, Reln | In3 | Interneuron 3 (Adarb2+) |
| 15 | | | Tle4, Zfpm2, Ldb2 | Glut1 | Glutamatergic 1 (Corticothalamic Neurons) |
| 16 | Gad2 | Adarb2, Nrxn3, Dlx6os1 | Reln | In3 | Interneuron 3 (Adarb2+) |
| 17 | Eomes | | Slc1a3, Sema3a, Slc17a6 | NPC | Neural progenitor cells |
| 18 | Neurod2, Neurod6, Rbfox1 | Ldb2 | Satb2, Dok5, Tle4 | Glut2 | Glutamatergic 2 (Callosal Projection Neurons) |
| 19 | Mki67 | Top2a | Zfpm2, Fabp7, Slc1a3 | RG | Radial Glial Cells |
| 20 | Mki67 | Top2a, Nrxn3, Dlx6os1 | | NSC | Neural Stem Cell |
| 21 | Gad2 | Isl1 | Nrxn3, Sema3a | In4 | Interneurons 4 (Isl1+) |
| 22 | Mki67 | Tnc, Top2a, Slc1a3 | Zfpm2 | RG | Radial Glial Cells |
| 23 | Rbfox3 | Slc1a3 | | Glut6 | Glutamatergic 6 (Unspecified Excitatory Neurons 2) |
| 24 | | Zfpm2 | Tle4 | Glut1 | Glutamatergic 1 (Corticothalamic Neurons) |
| 25 | Neurod2, Neurod6 | | Sema3a, Slc17a6 | Glut4 | Glutamatergic 4 (Migrating Pyramidal Neurons 2) |
| 26 | Gad2, Rbfox1 | Nrxn3 | Lhx6, Satb2, Sema3a | In1 | Interneuron 1 (Lhx6+/Sst -) |
| 27 | | | Olig1, Olig2, Pdgfra, Sox10, Zfpm2, Slc1a3 | OPC | Oligodendrocyte Progenitor Cells |
| 28 | Rbfox3 | Satb2, Dok5 | Ldb2 | Glut2 | Glutamatergic 2 (Callosal Projection Neurons) |
| 29 | | Slc1a3, Fabp7 | Zfpm2 | Astro | Astrocytes |
| 30 | | Reln | Slc17a6 | Neuron | Unspecified Neuron |
| 31 | Rbfox1, Rbfox3 | Tle4, Zfpm2 | Adarb2, Nrxn3 | Glut1 | Glutamatergic 1 (Corticothalamic Neurons) |
| 32 | | Col4a1, Col4a2 | Cldn5, Zfpm2 | Endo | Endothelial/Pericytes |
| 33 | | | C1qa, C1qb | Mg | Microglia |
| 34 | Rbfox1 | Satb2, Reln | Ldb2 | Glut5 | Glutamatergic 5 (Unspecified Excitatory Neurons 1) |
| 35 | Gad2 | Lhx6, Nrxn3 | | In1 | Interneuron 1 (Lhx6+/Sst -) |

**Table S3: Cluster Annotations by ATAC-seq Gene Activities Markers**

| Cluster | Class | Top 10 Subclass Markers | Other Subclass Markers | Annotation | Expanded Cell types |
|---|---|---|---|---|---|
| 1 | Neurod2, Neurod6 | | Satb2, Unc5d, Sema3c, Dok5 | Glut3 | Glutamatergic 3 (Migrating Pyramidal Neurons 1) |
| 2 | Neurod2, Neurod6 | | Tle4, Zfpm2 | Glut1 | Glutamatergic 1 (Corticothalamic Neurons) |
| 3 | Gad2, Rbfox3 | Lhx6 | Dlx6 | In1 | Interneuron 1 (Lhx6+/Sst -) |
| 4 | Neurod2, Neurod6 | Satb2 | Unc5d, Dok5 | Glut2 | Glutamatergic 2 (Callosal Projection Neurons) |
| 5 | Neurod2, Neurod6 | Satb2 | Dok5 | Glut2 | Glutamatergic 2 (Callosal Projection Neurons) |
| 6 | Neurod2, Rbfox1, Rbfox3 | | Sema3c | Glut4 | Glutamatergic 4 (Migrating Pyramidal Neurons 2) |
| 7 | Gad2, Rbfox1 | | Isl1, Drd2, Adarb2 | In4 | Interneuron 6 (Unspecified Interneuron) |
| 8 | Gad2 | Dlx6 | Adarb2 | In3 | Interneuron 3 (Adarb2+) |
| 9 | | Fabp7 | Col4a2, Slc1a3 | Astro | Astrocytes |
| 10 | Gad2, Rbfox1 | Drd2 | Tle4, Dlx6 | In5 | Interneurons (Drd2+) |
| 11 | Neurod2, Neurod6 | Satb2 | | Glut2 | Glutamatergic 2 (Callosal Projection Neurons) |
| 12 | Neurod6 | | Sema3c, Slc1a3 | Glut4 | Glutamatergic 4 (Migrating Pyramidal Neurons 2) |
| 13 | Neurod2, Neurod6, Rbfox1 | Satb2 | Unc5d, Sema3c | Glut2 | Glutamatergic 2 (Callosal Projection Neurons) |
| 14 | Gad2 | | Col4a1, Drd2 | In6 | Interneuron 6 (Unspecified Interneuron) |
| 15 | Neurod6 | | Tle4, Zfpm2, Sema3c | Glut1 | Glutamatergic 1 (Corticothalamic Neurons) |
| 16 | Gad2, Rbfox3 | Adarb2 | | In3 | Interneuron 3 (Adarb2+) |
| 17 | Eomes | | Unc5d, Col4a2, Slc1a3 | NPC | Neural progenitor cells |
| 18 | Neurod2, Neurod6 | | Satb2 | Glut2 | Glutamatergic 2 (Callosal Projection Neurons) |
| 19 | Eomes, Mki67 | Olig2 | Fabp7, Top2a, Slc1a3 | RG | Radial Glial Cells |
| 20 | Gad2, Mki67 | | Olig2, Top2a | NSC | Neural Stem Cell |
| 21 | Gad2, Rbfox1 | Isl1 | | In4 | Interneurons 4 (Isl1+) |
| 22 | Mki67 | | Olig2, Top2a | RG | Radial Glial Cells |
| 23 | | | | Unknown | No clear markers |
| 24 | | | Tle4, Zfpm2, Drd2 | Glut1 | Glutamatergic 1 (Corticothalamic Neurons) |
| 25 | Neurod2, Neurod6 | | | Glut5 | Glutamatergic 5 (Unspecified Excitatory Neurons) |
| 26 | Gad2 | Lhx6 | | In1 | Interneuron 1 (Lhx6+/Sst -) |
| 27 | Mki67 | | Fabp7, Olig2 | RG | Radial Glial Cells |
| 28 | Neurod6 | | Satb2 | Glut2 | Glutamatergic 2 (Callosal Projection Neurons) |
| 29 | | Fabp7 | Zfpm2, Col4a2, Olig2 | Astro | Astrocytes |
| 30 | | Reln | Col4a1, Drd2 | Neuron | Unspecified Neuron |
| 31 | | | Tle4, Zfpm2, Lhx6 | Glut1 | Glutamatergic 1 (Corticothalamic Neurons) |
| 32 | | | Col4a1, Col4a2 | Endo | Endothelial/Pericytes |
| 33 | Mki67 | C1qb | C1qa | Mg | Microglia |
| 34 | | | Unknown | Unknown | No clear markers |
| 35 | Gad2, Rbfox3 | Lhx6 | In1 | In1 | Interneuron 1 (Lhx6+/Sst -) |

**Table S4: Computational Programs**

**R programs**

| **Names** | **Version** | **Website (if applicable)** |
|---|---|---|
| ArchR | 1.0.2 | https://www.archrproject.com/ |
| BSgenome.Mmusculus.UCSC.mm10 | 1.4.2 | |
| ComplexHeatmap | 2.14.0 | https://jokergoo.github.io/ComplexHeatmap-reference/book/ |
| condiments | 1.6.0 | https://hectorrdb.github.io/condiments/articles/condiments.html |
| cowplot | 1.1.1 | |
| dplyr | 1.1.2 | |
| gbeeswarm | 0.7.2 | |
| gghighlight | | |
| ggplot2 | 3.4.2 | |
| ggpubr | 0.6.0 | |
| harmony | 0.1.1 | |
| patchwork | 1.1.2 | |
| pathfindR | 2.1.0 | https://egeulgen.github.io/pathfindR/index.html |
| perm | 1.0-0.2 | |
| R | 4.2.3-4.4 | |
| RColorBrewer | 1.1-3 | |
| readr | 2.1.4 | |
| rliger | 1.0.0 | |
| Rstudio | 0.14 | |
| Seurat | 4.9.9 | https://satijalab.org/seurat/ |
| Signac | 1.9.0 | https://stuartlab.org/signac/ |
| SingleCellExperiment | 1.20.1 | |
| slingshot | 2.6.0 | |
| stringr | 1.5.0 | |
| tibble | 3.2.1 | |
| tidyverse | 2.0.0 | |
| tradeseq | 1.12.0 | |

**Python**

| **Names** | **Version** | **Website (if applicable)** |
|---|---|---|
| MACS2 | | |
| velotcyto | 0.17.17 | https://velocyto.org/ |
| scVelo | 0.2.5 | https://scvelo.readthedocs.io/en/stable/index.html |
